# The Kinetics of Carbon-Carbon-Bond Formation in Metazoan Fatty Acid Synthase and its Impact on Product Fidelity

**DOI:** 10.1101/2024.07.03.601458

**Authors:** Christian Gusenda, Ana R. Calixto, Joana R. da Silva, Pedro A. Fernandes, Martin Grininger

**Affiliations:** Institute of Organic Chemistry and Chemical Biology, Buchmann Institute of Molecular Life Sciences, Goethe University Frankfurt, Max-von-Laue-Str. 15, 60438 Frankfurt am Main, Germany; LAQV, REQUIMTE, Departamento de Química e Bioquímica, Faculdade de Ciências, Universidade do Porto, Rua do Campo Alegre s/n, 4169-007 Porto, Portugal

## Abstract

Fatty acid synthase (FAS) multienzymes are responsible for de novo fatty acid biosynthesis and crucial in primary metabolism. Despite extensive research, the molecular details of the FAS catalytic mechanisms are still poorly understood. For example, the β-ketoacyl synthase (KS) catalyzes the fatty acid elongating carbon-carbon-bond formation, which is the key catalytic step in biosynthesis, but factors that determine the speed and accuracy of his reaction are still unclear. Here we report enzyme kinetics of the KS-mediated carbon-carbon bond formation, enabled by a continuous fluorometric activity assay. We observe that the KS kinetics are adapted to the length of the bound fatty acyl chain, and that the KS is also responsible for the fidelity of biosynthesis by preventing intermediates from undergoing KS-mediated elongation. To provide mechanistic insight into KS selectivity, we performed computational molecular dynamics (MD) simulations. Intriguingly, the KS protomers within the dimer exhibit positive cooperativity, investigated by mutational studies and acyl-carrier analysis, which likely serves the regulation of biosynthesis. Advancing our knowledge about the KS molecular mechanism will pave the ground for engineering FAS for biotechnology applications and the design of new therapeutics targeting the fatty acid metabolism.

## Introduction

Fatty acids are essential for life. They are signaling molecules, components of membrane-building phospholipids, esterified to triacylglycerols for energy storage, and stabilize or localize other molecules for cellular functions.^[1,2]^ Fatty acids are de novo biosynthesized by fatty acid synthases (FASs) by condensation and processing of the activated carbonic acids, acetyl- and malonyl-coenzyme A (CoA), to long-chain fatty acids, most commonly palmitic acid (C16) or stearic acid (C18).^[3]^ Advancing the knowledge of FASs is important due to the fundamental role they play in primary metabolism. In addition, FAS presents a promising target for treating a wide range of diseases, like obesity and cancer,^[4–9]^ and a better mechanistic understanding of the FAS molecular properties can prepare grounds for new therapeutic strategies.

While sharing the same biosynthetic principle (Figure 1A), FASs are divided by their overall structural organization into two main systems: type I and type II FASs. Type II FASs is mostly present in bacteria, plants and eukaryotic organelles of prokaryotic descent. This type consists of single, monofunctional proteins, which are regulated by discrete genes.^[10]^ In contrast to this, type I FASs are multi-enzyme proteins in which catalytic domains are covalently linked to each other (Figure 1B). The overall architecture of type I FAS in metazoa is identical in the related polyketide synthases (PKS), which additionally share the same reaction cascade but produce a higher variety of products.^[11–18]^ The concept of multi-enzyme complexes brings advantages such as high intermediate concentrations, which lead to high turnover rates, reduction of side reactions by direct substrate shuttling and coordinated gene regulation. The evolutionary fusion of discrete type II enzymes to domains in type I systems probably induced kinetic alterations that are not yet fully understood.

**Figure 1.**
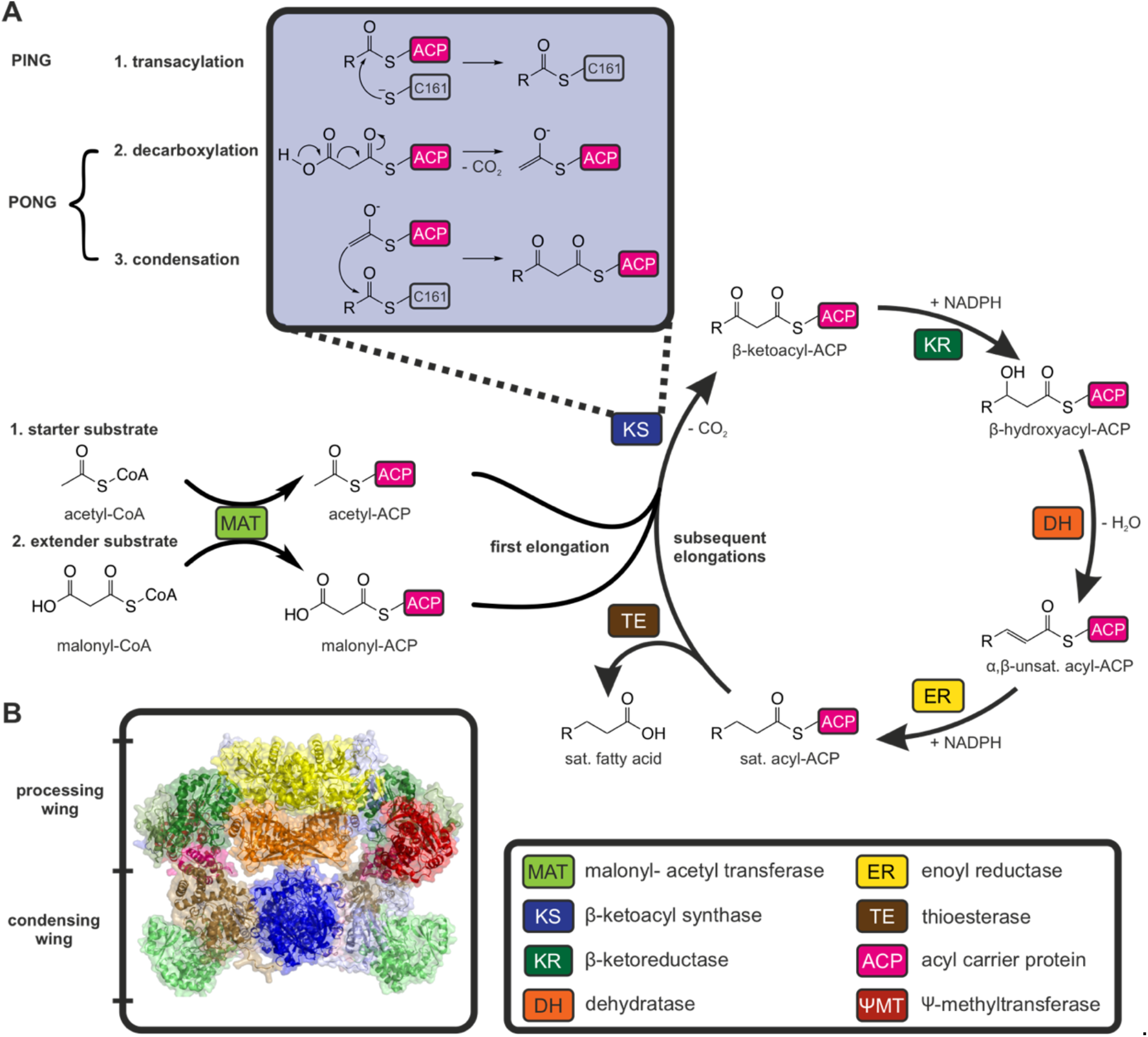
Mechanism of the fatty acid synthesis. **A** The acetyl moiety of acetyl-CoA is transferred to an acyl carrier protein (ACP) by a malonyl-acetyl transferase (MAT) catalyzed reaction, and subsequently loaded to the active side cysteine of the β-ketoacyl synthase (KS) (1. Transacylation). Free ACP is equipped with a malonyl moiety of the extender substrate malonyl-CoA. Malonyl is decarboxylated to an enolate (2. Decarboxylation), which acts a nucleophile in a Claisen-like condensation reaction with the KS bound acetyl (3. Condensation). The β-ketoacyl condensation product, that is bound to ACP, is transferred to the β-ketoreductase (KR), which reduces the keto group to a hydroxy group, to the dehydratase (DH) for dehydratation, and to the enoyl reductase (ER), which reduces the unsaturated acyl-ACP to the saturated compound. The cycle is repeated until a chain length of typically 16 carbon atoms is reached (palmitic acid). The thioesterase (TE) catalyzes the hydrolysis of the product from the ACP, which releases the free fatty acid. **B** Structure of murine fatty acid synthase from the alpha fold database (AFDB: AF-P19096-F1) with indication of condensing wing (KS, MAT) and processing wing (KR, DH, ER, Ψ MT).

The structure of the metazoan FAS was determined in 2006 with the X-ray structure of pig FAS that revealed an x-shaped domain architecture. Since then, structural insight has been further refined with additional X-ray, cryogenic electron microscopic data and molecular modelling.^[3,19–23]^ These findings, coupled with detailed biochemical studies, offer a detailed understanding of the iterative synthesis in metazoan FASs. Remarkably, metazoan FASs operate at extremely high rates, several-fold faster than related PKS. Aiming to understand the remarkable metazoan FAS catalytic efficiency, the KS domain deserves a closer look, particularly, as it catalyzes the pivotal carbon-carbon bond formation during fatty acid synthesis. The reaction mechanism of the KS follows a sequential ping-pong mechanism (Figure 2).^[4]^ The first step is the transacylation part of the reaction, which involves the transfer of an acyl from ACP to the active site residue Cys161 (murine FAS numbering is used throughout this article). The second half-reaction is the decarboxylation of the elongating malonyl residue and its Claisen-like ester condensation with the Cys161-bound acyl residue (Figure 1A). So far, only the kinetics of the transacylation and the decarboxylation part reactions of metazoan FAS have been studied.^[24–26]^ However, by examining portions of the KS catalysis, these studies were incapable to account for the features of the overall reaction. In addition, these studies missed valuable enzymatic details as e.g. the characterization of the domain-domain interaction during the KS catalyzed reactions, which has been suggested to be a key factor in orchestrating biosynthesis in type I and II fatty acid and polyketide biosynthesis.^[27–33]^ The condensation part of the reaction is hardly characterized at all, and so far, focused on bacterial type II KS. ^14^C-labelled malonyl-CoA was used in the first study of bacterial KS.^[34]^ Later, an enzyme-coupled assay enabled continuous monitoring of the full KS-catalyzed reaction. Here, stand-alone β-ketoacyl reductases are harnessed to detect product formation by spectral changes upon consumption of the cofactor NADPH (Figure 3A).^[35,36]^ Several more functional and structural studies were conducted on bacterial KS enzymes.^[37–41]^

**Figure 2.**
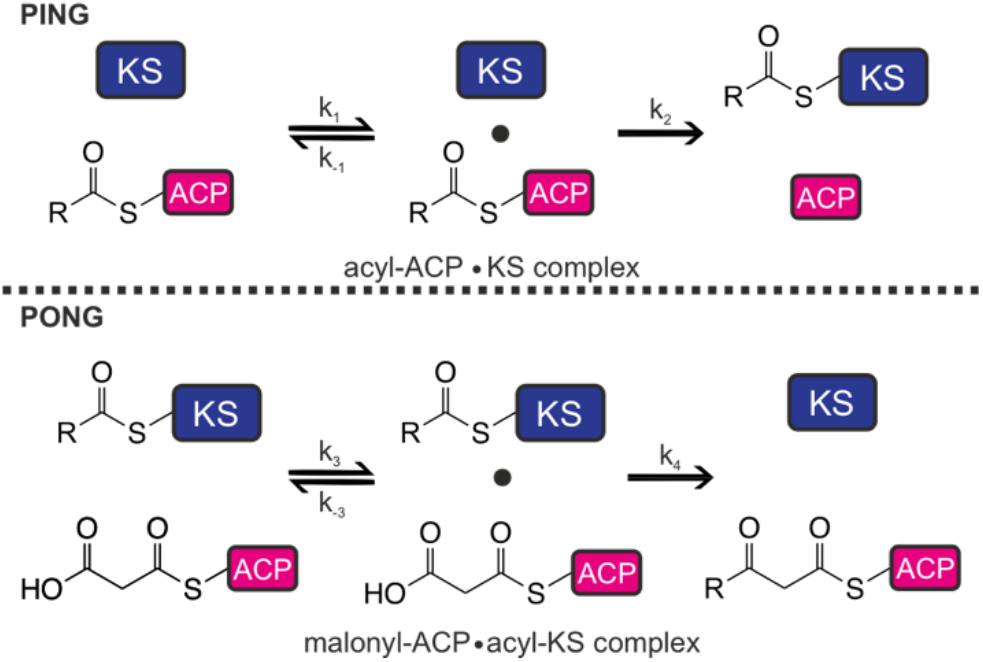
Overall reaction of KS-mediated chain elongation split in transacylation (ping step) and condensation (pong). During the ping step, an enzyme-substrate complex is formed and exists in equilibrium with the dissociated enzyme and substrate. The rate with which the complex forms is called k_1_. Rate k_-1_ describes the complex dissociation rate. The transacylation of the acyl moiety to the KS Cys161 and the subsequent dissociation of the enzyme-product complex happens with the rate k_2_. During the pong step, the enzyme-substrate complex of acyl KS and malonyl ACP is formed with the associating rate of k_3_ and dissociating rate of k_-3_. The decarboxylation, condensation reaction and dissociation of the enzyme-product complex are summarized by the rate k_4_.

**Figure 3.**
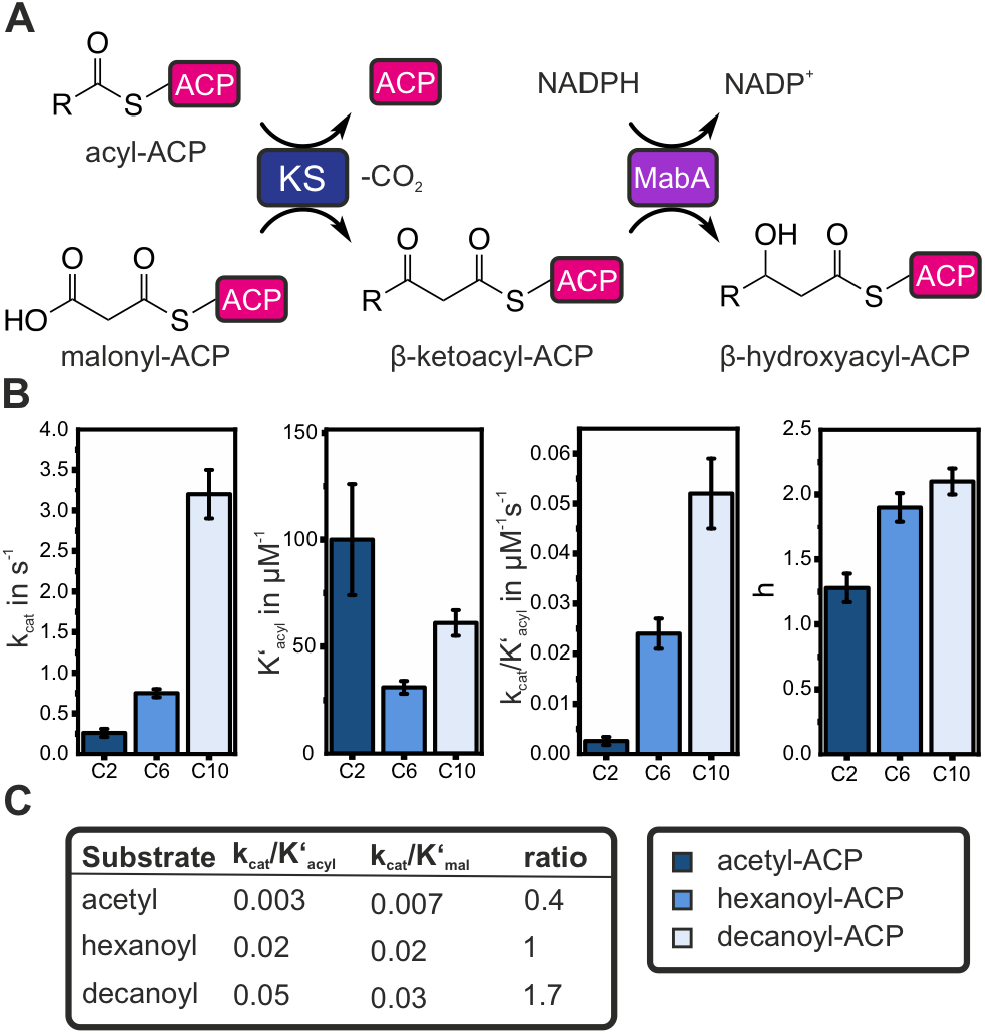
Kinetic analysis of the wildtype KS. A Scheme of the MabA-coupled assay. The MabA-coupled assay allows continuous monitoring of the entire KS-mediated two-step condensation reaction by the NADPH-dependent reduction of the condensation product. Maintaining an excess reduction capacity with high concentrations of reductase allows to indirectly monitor the formation of β-keto acyl-ACP and, therefore, the KS activity. The formation of the β-keto acyl product is proportional to the oxidation of the reducing agent NADPH and, therefore, to NADPH fluorescence quenching. B Turnover number k_cat_ of the wildtype KS for acetyl-ACP, hexanoyl-ACP and decanoyl-ACP; Michaelis-Menten constant derived value K’_acyl_ of Hill equation; Enzymatic efficiency k_cat_/K’_acyl_; Hill constant h. Bar heights represent the means of three biological replicates, error bars show standard deviations. All measurements were performed at 25°C. C Enzymatic efficiencies of the wildtype KS regarding different chain lengths of acyl substrates and corresponding efficiency regarding malonyl-ACP. Values are given in s^-1^µM^-1^.

In the present study, we asked how the KS-mediated condensation reaction affects the kinetics and specificity of cytoplasmatic de novo fatty acid synthesis. Specifically, we asked whether the condensation reaction contributes to the substrate specificity of the KS and which role the ACP-KS domain interactions plays. That is why we determined the full-reaction kinetics of the type I mFAS (metazoan FAS)-KS domain for ACP-bound substrates. To do so, we employed the enzyme-coupled assay to characterize the kinetics of the KS domain for three natural substrates, which revealed chain length specificity of the enzyme, as well as for non-natural substrates, which indicated that the KS is responsible for the fidelity of fatty acid biosynthesis. The experimental insight to substrate discrimination was complemented by molecular modelling. Compared to saturated fatty acids, the *in-silico* substrate molecular dynamics (MD) simulations revealed that the fatty acid cycle intermediates bind to the KS binding tunnel in catalytically unproductive positions. With the help of the enzyme-coupled assay, we further identified the cooperative working mode of the KS dimer. Although the molecular origin of the cooperative response remained elusive, we demonstrate that domain-domain interactions and the combination of starting and elongating substrates are relevant to the cooperative response

## Results and Discussion

### Enzyme Coupled Fluorometric Assay

We began this study with the hypothesis that the responsibility for maintaining high reaction rates while ensuring accuracy in terms of product chain length and chemistry predominantly lies with the KS domain by choosing appropriate substrates for condensation. To understand how the KS domain accomplishes this complex task, we initiated the analysis of the KS enzyme’s kinetic properties. We decided to work with the KS domain of the type I murine FAS because we had established recombinant access to this protein before. The sequence of murine FAS is overall 82% identical to human FAS, with 89% sequence identity for the KS domain. Since the KS domain is not accessible as an alone-standing protein in sufficient yield and quality in recombinant protein production, the KS-MAT didomain, carrying mutation S581A (functional MAT knockout, MAT^0^) and termed KS-MAT^S581A^ in the following, was used in this assay (Figure S1).^[42]^

The KS-mediated condensation reaction does not involve substrates or products that can easily be monitored continuously by spectroscopic means, nor does it include consumption or production of cofactors with spectroscopically accessible properties. When seeking for a direct and continuous assay as a platform for the thorough analysis of the KS kinetics, we therefore decided to adapt the *Mycobacterium tuberculosis*-based coupled assay for type I FAS (Figure 3A).^[35]^ The assay is based on the *in vitro* catalysis of the KS-mediated condensation reaction and adds a subsequent reduction step from fatty acid biosynthesis by including a NADPH-based β-ketoacyl reductase. The reduction and hence the prior condensation can be monitored by detecting fluorescence quenching of NADPH.

The use of acyl-ACP substrates beyond the chain length of ten carbon atoms was limited due to their decreased half-lives (Figure S10). We sought to switch to (S, N)-acetylcysteamine (SNAC)-derived model substrates instead of ACP tethered substrates to circumvent acyl-ACP stability problems. However, due to poor solubility at acyl chain lengths larger than six carbon atoms, these substrates proved to be unsuitable for our needs. Coenzyme A tethered substrates, on the other hand, induce substrate inhibition at low concentrations such that they are unsuitable for achieving an understanding of the KS that closely mirrors natural conditions (Figure S8). In conclusion, we conducted this study with acyl-ACPs up to an acyl chain length of C10.

### Determination of Kinetic Constants

The enzyme specific kinetic constants K_m_ and k_cat_ are commonly used to compare the enzymatic efficiency (k_cat_/K_m_) between different enzyme variants or different substrates. To determine the kinetic constants of the overall two-step condensation reaction, a starter acyl substrate and an elongation malonyl substrate were applied in different concentrations to measure the initial reaction velocity with the MabA assay (Figure S4). The obtained data (initial reaction velocity against acyl-ACP concentration) exhibited a sigmoidal shape, which indicates a cooperative behavior of the KS. To account for the deviation from a simple hyperbolic saturation curve, data was globally fitted using the Hill equation to obtain kinetic values (equation 1). Value v_max_ reflects the maximal velocity of the enzyme. K’ is related to the Michalis-Menten constant K_m_, which reflects the substrate concentration at 0.5 x v_max_, but additionally K’ takes substrate occupancy and affinity into account. Hill constant h is a measure of cooperativity in enzyme catalysis, where binding of substrate S to the first active site enhances (or decreases) the affinity of the same substrate S to other active sites. The sigmoidal response was previously observed for the transacylation half reaction of mFAS-KS as well as for type II homologs.^[24,43]^ Further, we noticed that the steepness of the sigmoidal titration curve depends on the chain length of the substrates indicating that cooperativity is a function of acyl chain length (Figure 3B).

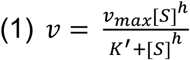

The enzymatic efficiency k_cat_/K^’^ can be calculated for both the ping and the pong step of a double displacement reaction (Figure 2, equation 2).^[44]^ The catalytic efficiency of the ping step is, therefore, reflected by k_cat_/K^’^_acyl_, whereas the efficiency of the pong step is reflected by k_cat_/K^’^_mal_.

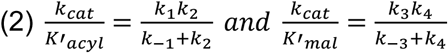

As depicted in Figure 3C, enzymatic efficiencies of KS increase with chain length in both the ping and pong steps. According to equation 2, an increased k_cat_/K^’^ can have various origins, including **a**) an increased KS affinity towards the substrate and, therefore, an equilibrium shift towards the enzyme-substrate complex (increased k_1_ or k_3_ over k_-1_ or k_-3_ respectively), **b**) an increased reaction rate (increased k_2_ or k_4_) or **c**) an increased dissociation of the enzyme-product complex (increased k_2_ or k_4_).

In the case of the ping step, a change in the dissociation of the enzyme-product complex (**c**) is unlikely, as, after delivering the substrate to the KS, the unloaded ACP will most likely have the same affinity towards the acyl KS, regardless of the identity of the previously bound substrate. Therefore, the increase in the enzymatic efficiency of the ping step with acyl chain length will be caused by either the affinity of the acyl ACP towards the KS (**a**) or the transacylation rate (**b**). In case of the pong step, a change in affinity towards malonyl ACP (**a**) or the decarboxylation rate of the same (**b**) is unexpected, given that the acyl chain is resting in the KS domain where it presumably does not play a significant role in these processes. Therefore, the increase in the enzymatic efficiency of the pong step with chain length will be caused by the condensation rate (**b**) or the dissociation of the enzyme-product complex (**c**). It is expected that the acyl chain lengths would have a more significant influence on the ping step than on the pong step of the KS mediated reactions. This assumption is well reflected by the more extensive shift of the ping enzymatic efficiency depending on the acyl chain length (Figure 3C). We conclude that the ping step, the transacylation reaction that delivers the acyl moiety to the KS, is the determining factor in the (chain length) specificity of the KS. In contrast, the pong step was identified to be the determining factor of specificity in trans-AT PKS.^[45]^

Our data reveals, that the enzymatic efficiency k_cat_/K^’^_acyl_ increases with chain lengths from two to ten carbon atoms (2.6 ± 0.8 mM^-1^s^-1^ for C2, 24 ± 3 mM^-1^s^-1^ for C6 and 53 ± 6 mM^-1^s^-1^ for C10; Figure 3B). Increasing enzyme activity with increasing chain length from very short to medium acyl chains for the KS catalyzed transacylation reaction was observed before when employing acyl-CoA substrates and monitoring CoA release.^[24,25]^ The enzymatic efficiency k_cat_/K^’^_acyl_ determined here is 1-2 orders of magnitude larger than determined by the transacylation assay previously, which we assume to originate from employing acyl-ACPs as the native substrates. However, despite the diverting absolute values, trends in the transacylation part-reaction and the overall KS mediated reaction are identical substantiating the finding that substrate specificity and cooperativity originate from the ping step.

The enzymatic efficiency of the mFAS-KS is in a comparable range to that of the homologous type II proteins KasA and KasB of *M. tuberculosis* and FabF and FabB of *E. coli* (Figure S5).^[35,46]^ However, the turnover number k_cat_ as well as the constant K_m,acyl_ are much smaller for these bacterial enzymes compared to the constants of the mFAS-KS. We suspect the large K’_acyl_ of mFAS-KS is a result of the high local concentration of the substrate due to its covalent linkage to the ACP domain. The high K_m_ ensures transient binding of the ACP to the KS, which is necessary to facilitate rapid docking to all domains in the FAS complex.

The turnover number of the full-length mFAS, measured by NADPH consumption of the internal keto reductase and enoyl reductase domains was previously determined to 3.1 ± 0.3 s^-1^ (Figure S5).^[42]^ In approximation, this overall turnover number corresponds to the average of turnover numbers of all chain lengths. We have recorded similar rates for KS with substrate C10-ACP only (3.2 ± 0.2 s^-1^; rates for other chain lengths are slower), such that the KS assay seems to underestimate KS condensation rates. Differences between the KS rates in the native enzyme complex (ACP integrated in the polypeptide) and the assay set up (stand-alone ACP) might origin in factors such as molecular steering. Besides that, the MabA assay accurately reflects the kinetics of the integrated KS.

### Analysis of the Acyl Binding as Origin of Cooperativity

As discussed in the preceding section, KS activity shows a sigmoidal but not simple Michaelis-Menten-type hyperbolic dependence on acyl-ACP concentration. The sigmoidal shape indicates a positive cooperative working mode of the dimeric KS domain, which essentially means that substrate turnover by one protomer promotes turnover by the other protomer (Figure 4A). For cooperativity, information about the catalytic status of protomers needs to be exchanged. In a previous work, a hydrogen bond network was proposed to enable protomer communication (Figure 4B-C):^[24]^ Upon comparing crystal structures of the KS-MAT didomain in the unbound (apo) and octanoyl-bound states, a conformational change in residues decorating the active site was overserved. Morphing these states suggests that during octanoyl binding the gatekeeping residue F395.A (chain A) rotates and induces a loop to shift in the other protomer harboring residues M132.B, Q136.B and M139.B (chain B). The loop displacement causes a conformational change of R137.B that is in interaction with binding site residues D158.B and A160.B through a hydrogen bond network (Figure 4C). To evaluate whether these coordinated changes are the molecular basis for cooperativity, amino acids of this hydrogen bond network were mutated. The steepest sigmoidal response curve of the wildtype was measured with C10-ACP; hence, mutants were evaluated by titration of C10-ACP (Figure S7). For mutants D158N, D158S and A160V, activity was abolished. As the protein quality seemed to be intact given the maintained dimeric state (Figure S6), the reasons for inactivity may be the inability of substrate binding or processing. Mutants R137K and R137A do not seem to alter the apparent Hill coefficient (3.2 and 3.0, respectively, compared to 2.9 of wildtype), which indicates that the residue R137 is not participating in protomer communication. Mutant A160G, however, lowered the apparent Hill coefficient (1.8) slightly, which we interpret as A160 being involved in cooperativity (Figure 4D).

**Figure 4.**
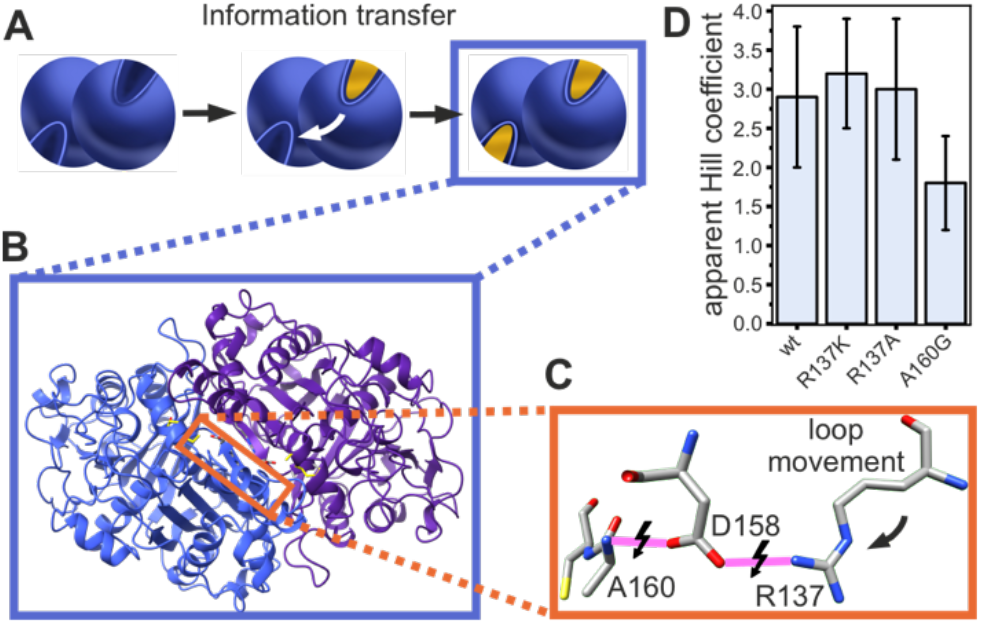
Proposed hydrogen bond network. **A** A simplistic scheme illustrating the principle of classical positive cooperativity. The binding of a substrate to one active site induces an affinity change of the second active site towards the same substrate. **B** Apparent Hill coefficient of the active mutants. Bars heights represent the means of three biological replicates and the black error bars standard deviation. **C** The KS structure in cartoon display shows the position of the hydrogen bond network (marked in orange). The two protomers are colored in blue and violet, respectively, and the bound substrate octanoyl is colored in yellow. **D** The hydrogen bond network of R137, D158 and A160. Hydrogen bonds are colored in pink and may change depending on binding site occupancy.

### Analysis of carrier binding as the origin of cooperativity

The apparent Hill coefficients of the wildtype and the R137 mutants are higher than expected for a dimeric protein, which should exhibit a Hill coefficient of two or lower. This indicates that the interpretation of the sigmoidal-shaped kinetics of the KS-mediated reaction as positively cooperative only taking the acyl moiety of the starting substrate into account does not capture the full complexity of the condensation reaction. In seeking to reveal the role of the substrate for KS cooperativity, we performed the coupled assay with substrates that are systematically truncated in the carrier part and analyzed whether sigmoidal profiles in velocity vs. substrate concentration plots were retained (Figure A). In nature, the KS substrates are tethered to the thiol moiety of a post-translationally introduced phosphopantetheine (Ppant) arm of ACP (Figure 1). Truncated versions of acyl-ACP are acyl-CoA and acyl-SNAC (Figure 5B): The larger CoA captures the interaction of Ppant with the binding tunnel that may facilitate correct binding of the substrates. We note that coenzyme A contains an additional adenosine-3’,5’-diphosphate, not present in acyl-ACP, which may induce additional attractive or repulsive interactions. The SNAC moiety is a mimic of the tip of the Ppant.

**Figure 5.**
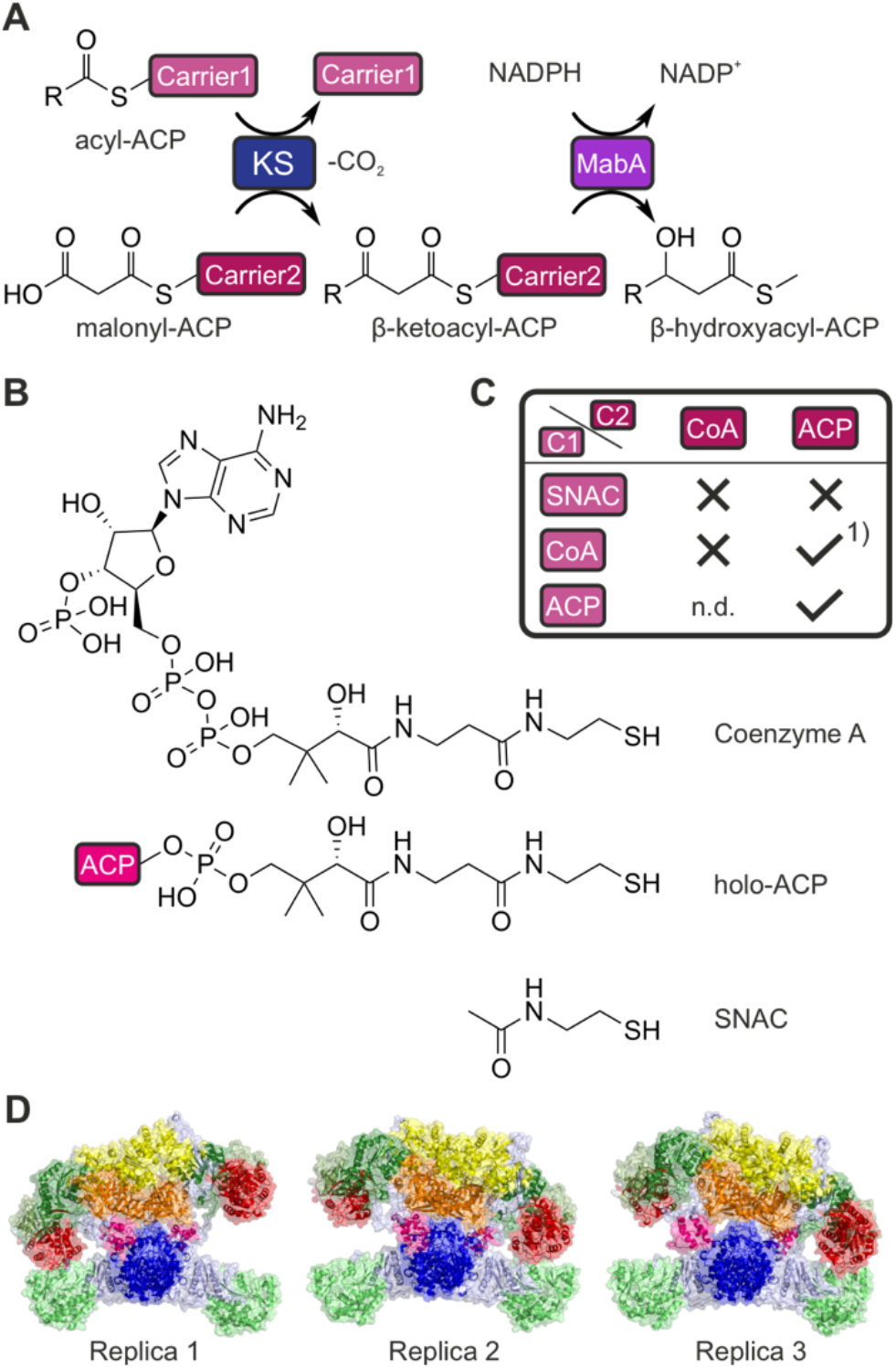
Analysis of the impact of the carrier in cooperativity. **A** Scheme of the MabA assay with two different carriers. The acyl-starter substrate is bound to carrier 1 and the malonyl extender substrate is bound to carrier 2. **B** In the MabA assay, various carrier types can be applied, including N-acetylcysteamine (SNAC), Coenzyme A (CoA) and ACP. All these carrier types share the N-acetylcysteamine group, whereas the CoA and ACP carriers additionally share the Ppant moiety. **C** Combination of carrier of starter and extender substrates changes KS kinetics. In the table, positive cooperativity is marked with a tick, whereas the absence of a sigmoidal response is marked with an X. 1) The analysis was performed with holo-ACP as the second substrate during transacylation measurement by Rittner et al.^[24]^ **D** The rat fatty acid synthase (rFAS) structure after the first, second and third 400 ns MD replicas. The simulation time of replica n. 1 was extended to 1 μs afterward. A rigid-body rearrangement of the two wings in rFAS promoted a conformation with slightly shorter and longer linkers in all MD simulations. The two ACP domains remained bound to the KS domains during the whole MD simulations in two of the three replicas. On the third replica, the ACP domains dissociated from the KS domains after a conformational shift. This behavior is expected and aligns with the well-known transient nature of the ACP-KS association. The ACP-KS interface was constituted by residues 45-49, 198-205, 297-298 (KS), and 47-74 (ACP), with an area of 197 Å^2^.

We used those carriers in the MabA assay, both as part of starter-carrier (donor) and extender-carrier (acceptor) substrates. Based on three datasets, we found that cooperativity does not simply stem from the acyl moiety, but its origin is more complex: Dataset 1, published previously by our lab: The use of hexanoyl-CoA (C6-CoA) as the donor substrate and holo-ACP as the acceptor substrate in the transacylation assay results in a sigmoidal shape of the collected data.^[24]^ Dataset 2: When C6-SNAC instead of C6-CoA (dataset 1) is used in the MabA assay (paired with the extender substrate Mal-ACP), the sigmoidal shape is lost (Figure 5C, S8). Dataset 3: The sigmoidal shape is also lost when the MabA assay is performed with C6-CoA and Mal-CoA as the extender substrate. From datasets 1 and 2, it can be concluded that cooperativity emerges already in the transacylation reaction and that cooperativity requires additional interactions provided by the Ppant arm (SNAC alone is not sufficient to induce cooperativity). Dataset 3 indicates that the general presence of ACP contributes to cooperativity.

Our findings are in accordance with the transacylation analysis of rat FAS, which utilized CoA and pantetheine as carrier and did not show any signs of cooperativity.^[25]^ It is further noted that the impact of (Mal-)ACP on the cooperative response was already seen in fluorometric binding assays of the KS FabF from *E. coli*.^[47]^

Since the structural prerequisite for cooperativity in the KS mediated reactions is the simultaneous binding of ACP domains, we next asked whether the mFAS scaffold inherently allows the binding of the ACP domains to both protomers of the KS dimer simultaneously. We performed molecular docking and molecular dynamics (MD) simulations, which confirmed that the linker of 10 amino acids (V2104-H2113), through which ACP is attached to KR, ensures sufficient conformational flexibility for simultaneous binding to both KS dimers (Figure 5D). The simulations further show a rigid-body conformational variability of the condensing and modifying wing leading to different distances of the KR and KS domains (Table S1), while no considerable changes in domain structures were observed (Figure S12-14). This overall conformation, which has the ACP docked at the KS in both protomers and a reduced angle between the two wings of the mFAS, serves as a picture of the mFAS-complex during the KS-catalyzed reaction in fatty acid synthesis. We would like to point out that previous data on heterodimeric mFAS, in which only one of two KS-domains was knocked out in function by mutation, can be captured via the cooperative properties: Here, the transacylation activity of the KS domain recorded with a non-cooperative (non-ACP) carrier reached expected 50%, while full fatty acid synthesis (ACP as carrier) did not reach 50% activity.^[48]^ These results further imply that the cooperativity of the KS dimer has an influence on the native fatty acid synthesis in the mFAS-complex.

### Monitoring Intermediates of the Fatty Acid Cycle

The metazoan FAS produces saturated fatty acids in high purity, which indicates that the selectivity of the KS for saturated acyl-moieties, as received after the ER-mediated reduction, is high. However, quantitative enzyme kinetic data on the selectivity of the condensation reaction is elusive, to the best of our knowledge. With the MabA-assay in hand, we sought to analyze the selectivity of the KS, particularly for preferring fully reduced substrates over intermediates of the fatty acid cycle. Intriguingly, previous work has shown that the mFAS is able to produce triacetic acid lactone (TAL) under non-reducing conditions at condensation rates of 16% of fatty acid production, which proves that the KS is, in general, able to condense β-keto acyl-units.^[42]^ To analyze KS selectivity, the fatty acid cycle intermediates (*R*)-β-hydroxybutyryl-ACP ((*R*)-HB-ACP) and crotonyl-ACP were applied in the MabA assay (Figure 6). The β-ketoacyl intermediate, the product of the KS-mediated elongation, is a substrate of MabA and was therefore not part of this study.

**Figure 6.**
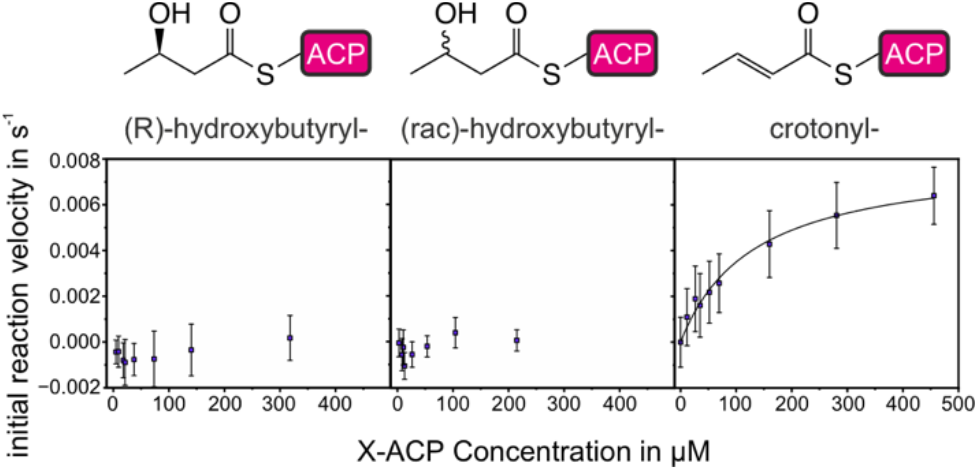
Kinetics of intermediates of the fatty acid cycle. Titration of (R)-HB-ACP, (rac)-HB-ACP and crotonyl-ACP. Measurements were performed with 1.5 µM KS and 70 µM Mal-ACP. All data points represent the means of three biological replicates, and the error bars represent the standard deviations.

Conducting the assay with (*R*)-HB-ACP, a representative second intermediate, did not result in detectable activity (Figure 6), suggesting that the KS is unable to condense β-hydroxyacyl-ACP and Mal-ACP. To rule out a false negative result caused by MabA failing to reduce the condensation product in the coupled reaction, the reaction mixture was additionally analyzed by urea PAGE (Figure S9). In case the KS accepts (*R*)-HB-ACP for condensation with Mal-ACP, both substrates would be consumed during the time course of the reaction. Urea-PAGE shows that the amounts of Mal-ACP and HB-ACP remain constant over one hour of reaction, confirming that (*R*)-HB-ACP is not a substrate for KS. We also questioned whether the KS may just counterselect against elongation of the (*R*)-HB-ACP enantiomer, which natively occurs as an intermediate in the fatty acid cycle but not against (*S*)-HB-ACP. However, when applying the racemic HB-ACP to the MabA assay, we could not observe any activity either (Figure 6).

For crotonyl-ACP, the third fatty acid cycle intermediate, we observed KS activity, which was lower compared to saturated fatty acids (k_cat_ ^app^ = 81 ± 5 x10^-4^ s^-1^ and K_m_ ^app^ = 0.13 ± 0.2 mM at 70 µM Mal-ACP). Thus, the discrimination of the FAS against unsaturated fatty acyl moieties (Figure 6) is less pronounced than for the β-hydroxy compounds. The specificity of the KS towards acetyl-ACP compared to crotonyl-ACP is k_cat_/K’_acetyl_ / k_cat_/K_m,crot_ = 50. The corresponding transition state energy difference Δ Δ G = −9.7 kJ/mol equals less energy than a weak hydrogen bond.

In conclusion, this data shows that the KS gatekeeps fatty acid synthesis by selecting against fatty acid cycle intermediates, thereby granting product fidelity. Data indicates that the discrimination of KS against the intermediates of the fatty acid cycle arises due to the different functionalization of the acyl moiety, with a double bond (assay substrate crotonyl-ACP) allowing less pronounced discrimination than a hydroxyl group (assay substrate HB-ACP).

### Computational analysis of substrate selectivity by KS

Our experimental data shows that the transacylation (ping) step can account for substrate specificity of the KS-mediated reaction (ping-pong, Figure 1A), which aligns with previous data monitoring the isolated transacylation kinetics.^[24]^ To obtain an atomic-level understanding of the specificity of the reaction, we performed MD simulations on the KS:acyl-Ppant complex formed during transacylation. Specifically, we compared the stability of acetyl, hexanoyl, decanoyl, crotonyl, and (R)-hydroxybutyryl substrates within the active site of the KS (Figure 7). The RMSD analysis of the protein backbone revealed the overall stability of KS throughout the 100 ns simulations (Figure S15). An analysis of the interatomic distances that are highly correlated with enzyme-substrate reactivity was carried out for each simulation (Table 1).

**Table 1.**
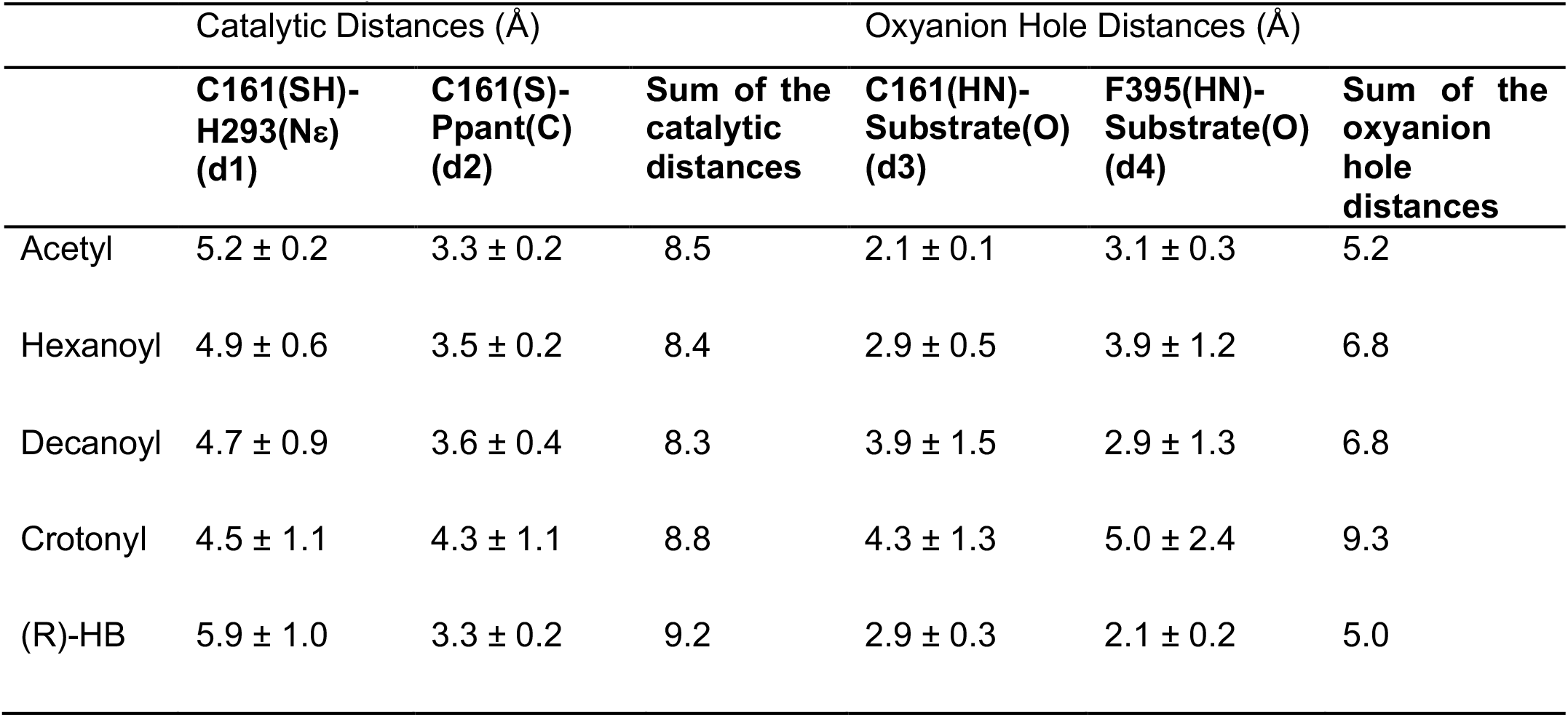
Average distances and standard deviations for important catalytic interactions to the first step of the reaction mechanism. All distances are presented in Ångström. Additionally, the sum of catalytic and oxyanion hole distances was computed to facilitate its interpretation.

|  | Catalytic Distances (Å) |  |  | Oxyanion Hole Distances (Å) |  |  |
| --- | --- | --- | --- | --- | --- | --- |
|  | C161(SH)-<br>H293(Nε)<br>(d1) | C161(S)-<br>Ppant(C)<br>(d2) | Sum of the<br>catalytic<br>distances | C161(HN)-<br>Substrate(O)<br>(d3) | F395(HN)-<br>Substrate(O)<br>(d4) | Sum of the<br>oxyanion<br>hole<br>distances |
| Acetyl | 5.2 ± 0.2 | 3.3 ± 0.2 | 8.5 | 2.1 ± 0.1 | 3.1 ± 0.3 | 5.2 |
| Hexanoyl | 4.9 ± 0.6 | 3.5 ± 0.2 | 8.4 | 2.9 ± 0.5 | 3.9 ± 1.2 | 6.8 |
| Decanoyl | 4.7 ± 0.9 | 3.6 ± 0.4 | 8.3 | 3.9 ± 1.5 | 2.9 ± 1.3 | 6.8 |
| Crotonyl | 4.5 ± 1.1 | 4.3 ± 1.1 | 8.8 | 4.3 ± 1.3 | 5.0 ± 2.4 | 9.3 |
| (R)-HB | 5.9 ± 1.0 | 3.3 ± 0.2 | 9.2 | 2.9 ± 0.3 | 2.1 ± 0.2 | 5.0 |

**Figure 7.**
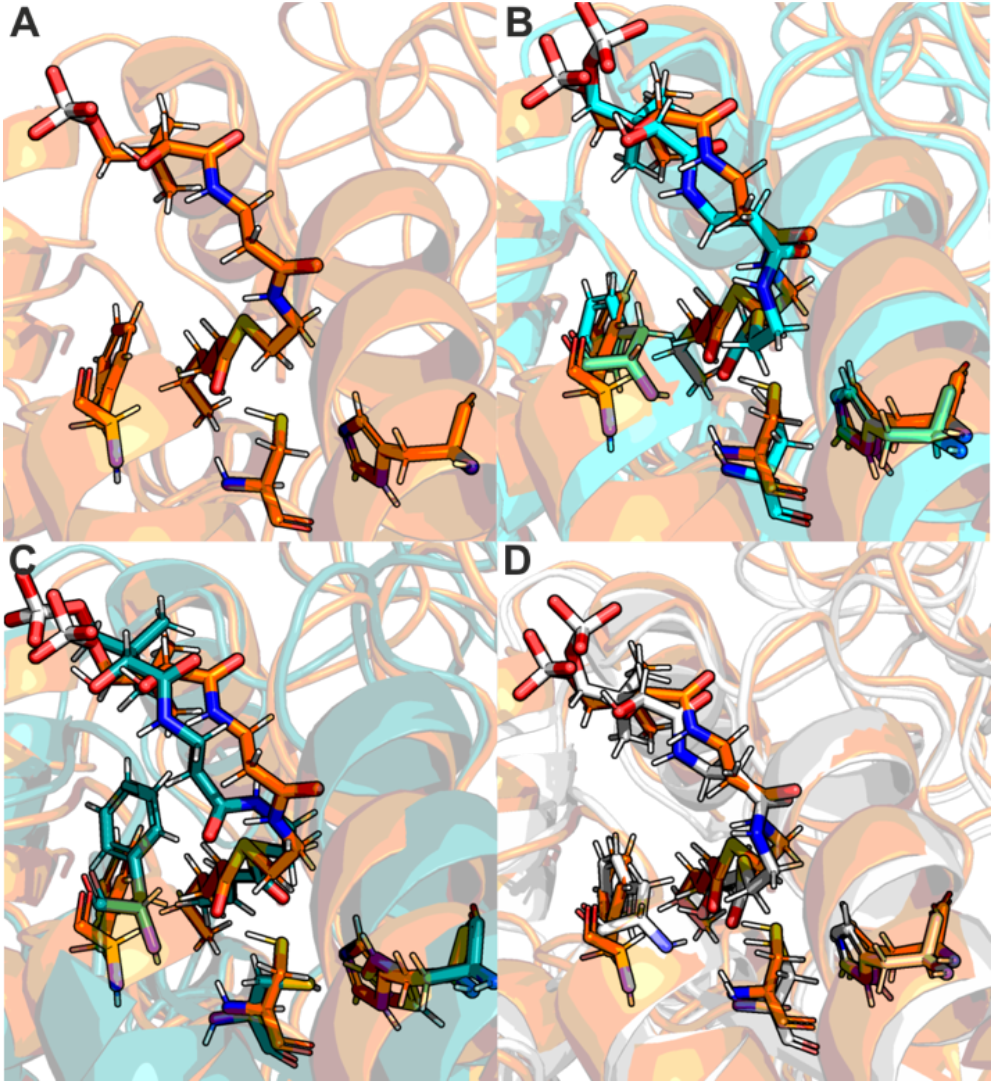
Snapshots from the MD Simulation of acyl-Ppant substrates in the KS active site. **A** Structure of hexanoyl-Ppant and the catalytic relevant residues C161, H293, F395. **B** Superposition of hexanoyl-Ppant (orange) and acetyl-Ppant (cyan). **C** Superposition of hexanoyl-Ppant (orange) and crotonyl-Ppant (teal). **D** Superposition of hexanoyl-Ppant (orange) and hydroxybutyryl-Ppant (silver).

For reference, the first step of the catalytic mechanism consists of Cys161 deprotonation by the N_ε_ atom of His293 and, in a probably concerted step, a nucleophilic attack on the Ppant-substrate thioester carbon by the Cys161 thiolate. However, we note that the distance to His293-N_ε_ might not be decisive because it has been shown earlier that a His293A mutant did not diminish transacylation activity.^[26]^ Additionally, an oxyanion hole composed of Phe395 and Cys161 establishes hydrogen bonds with the negative Ppant thiolate generated at the reaction transition state and stabilizes the transition state. The latter effect lowers the activation free energy of 18.8 kJ/mol at most.^[49]^ The distances of the four described interactions were analyzed (Figure S17).

Acetyl, hexanoyl, and decanoyl display low distances between catalytic residues (Table 1, Figure 7A-B), except for the Cys161-SH---His293-N_ε_ distance in acetyl-Ppant, known to be the least reactive of the three. Shorter oxyanion hole hydrogen bond distances partially compensate for this for the latter. This data indicates that the reactivity of the transacylation complex affects the enzymatic efficiency and, thus, specificity. However, data cannot rule out that the affinity of acyl-Ppant to KS is increased (an increase of k_1_ over k_-1_, see equation 2) and contributes to raising enzymatic efficiency.

Upon examination of the interatomic distances, it became evident that crotonyl displays higher, less favorable distances between catalytic residues and longer oxyanion hole hydrogen bonds (Figure 7C), indicating its inherent lower reactivity within the active site of KS.

R-HB, on the other hand, seems to be suitably aligned in the active site, given its close distances to Cys161-SH and to the oxyanion hole forming backbone amide groups (Table 1, Figure 7D). Still, the distance of the hydrogen bond between the Cys161-SH and the His293-N_ε_ is high, precluding the Cys161 deprotonation and subsequent nucleophilic attack on the Ppant (Table 1).

In general, the study showed that the experimentally evaluated enzymatic efficiencies of all substrates correlate well with the reactivity measured by interatomic distances of the catalytic residues. In addition, the study suggests that the transacylation kinetic is mostly correlated with the easiness of Cys161 deprotonation and nucleophilic attack on Ppant thioester carbon and, on a lesser scale, the stabilization of the negative charge accumulated at the Ppant thiolate by the oxyanion hydrogen bonds.

## Conclusion

The FAS multienzymes are crucial for the synthesis of long-chain fatty acids from acetyl-CoA and malonyl-CoA through a series of enzymatic reactions. Herein, the KS domain catalyzes the central carbon-carbon bond formation and thus holds key responsibility for the fidelity of long-chain fatty acid synthesis. So far, the enzyme kinetic analysis of mFAS catalytic domains were limited by using substrate analogs acyl-Ppant or acyl-CoA, resulting in properties of the reactions not being observable.^[25,50,51]^ Here, for the first time, we paint the full picture of the KS kinetics using an enzyme-coupled assay with close-to-nature kinetics. Our analysis reveals that the KS catalysis is dependent on the chain length of the fatty acid intermediate. Specifically, it exhibits increasing substrate specificity from short to medium saturated fatty acids, which unveils that the first cycle (condensation of acetyl-and malonyl moieties) is the slowest in fatty acid synthesis.

The KS kinetic properties are particularly interesting when considered in the context of the MAT domain, which loads and unloads ACP domains with acyl groups (Figure 1A). The relatively slow elongation of short acyl chains by the KS is in contrast with the very rapid transacylation of these compounds by the MAT. For biosynthesis, this means that short acyl chains bound to FAS will exhibit high tendency to be unloaded by the MAT. In this function, the MAT depends on the concentration of acetyl-CoA and malonyl-CoA as substrates, as they compete for the MAT binding pocket. The concentration of malonyl-CoA has been described to vary depending on the metabolic state of a cell,^[52]^ such that offloading of short acyl chains will be promoted at low concentrations, while at higher malonyl-CoA concentrations FAS will overcome the kinetic lag phase imposed by KS. Thus, via the interplay of KS and MAT, FAS is offered direct feedback on the metabolic state of the cell.

The titration of substrates revealed a positive cooperative response in KS activity, which was further investigated. We determined Hill coefficients for substrates with different chain lengths and bound to different carriers (ACP, CoA and SNAC), as well as of KS mutants, which vary a hydrogen bond network that bridges the protomers of the KS dimer. Our data suggests that the ACP plays a crucial role in triggering the cooperative response of the KS. The molecular basis for cooperativity was not fully elucidated, and the impact it has on biosynthesis remains unclear. However, we speculate that cooperativity, which increases from short to medium chain length, contributes to increasing enzymatic efficiency of the mFAS by synchronizing the condensation steps in the two reactions clefts and confining the higher-order conformational dynamics of mFAS, that is the positional variability of the domains within the multidomain complex.

Further, this study showed that the fidelity of the mFAS derives from the discrimination of the KS against fatty acid cycle intermediates (tested with HB-ACP and crotonyl-ACP). This discrimination was demonstrated with the enzyme-coupled assay. MD simulations showed elongated, unproductive catalytic distances of the crotonyl-Ppant and HB-Ppant complexes compared to that of saturated intermediates. The selectivity of the reaction of the KS emerges from the first step (ping step) of the double replacement reaction mechanism. This is plausible, as the sequence of steps in KS-mediated condensation is reversible up to the loading into the KS (before decarboxylating Claisen-like ester condensation), thus allowing correction of false KS loading without incurring energetic costs.

As shown by the kinetic measurements, the presented MabA assay is suitable for monitoring reaction kinetics and enzyme-specific properties. As the KS is the most conserved domain in the genetically closely related megasynthases FAS and PKS across species, it is likely that the same method can be applied to KS from several other systems, including PKSs. The presented study will contribute to a deeper understanding of the reaction sequence, domain interactions and specificities in type I multienzymes. It will facilitate protein engineering approaches that, in turn, can grant access to new-to-nature biosynthetic pathways. In addition, the details about KS catalysis can provide valuable insights for the design of new FAS-inhibition therapeutics for the treatment of obesity and cancer.

## Supporting Information

The authors have cited additional references within the Supporting Information. ^[53–64]^

## Acknowledgements

We thank A. Rittner who started the project with us. We thank E. Helfrich and his lab for support in using LC-MS instrumentation, and Maria Joao Ramos for supervision and analysis of MD simulations. Support for this work was received from Verband der Chemischen Industrie with a Kekulé Scholarship awarded to Christian Gusenda, as well as from the German Research Foundation (DFG grant GR3854/10-1 to MG). Pedro Fernandes and Ana Calixto acknowledge the Laboratório Associado para a Química Verde (LAQV), which is financed by FCT/MCTES within the scope of projects LA/P/0008/2020 UIDP/50006/2020, and UIDB/50006/2020.

## Notes

### Competing Interest Statement

The authors have declared no competing interest.

