## Supplementary Information for "The Kinetics of Carbon-Carbon-Bond Formation in Metazoan Fatty Acid Synthase and its Impact on Product Fidelity"

### Supporting Information

#### Additional Materials and Methods

**Cloning.** Plasmids of MabA, Sfp, ACP and KS-MAT were readily available as stated in the section “sequences” of this SI. The point mutations for cooperativity studies were introduced pcr based with primers mentioned in the section “primers”. CloneAmp$^{TM}$ HiFi PCR Premix (Takara) was used. The initial denaturation of the template KS-MAT$^{S581A}$ was run at 98°C for 180 s, followed by 23 cycles of 98°C for 10 s, 75°C for 20 s and 72°C for 90 s. The final elongation was performed at 72°C for 420 s. The template was digested with DpnI (NEB) at 37°C for 1 h. The PCR products were purified via agarose gel electrophoresis and extracted using the NucleoSpin® Clean-Up Kit (Macherey-Nagel). Purified DNA was transformed unto Stellar$^{TM}$ competent cells (Takara) and grown in LB-Agar (containing 100 µg/mL Amp and 1% glucose). One clone each was used to inoculate 8 mL of LB medium, which was incubated at 200 rpm and 37°C in culture tubes O/N. The plasmid was extracted from the cells using GeneJET Plasmid Miniprep Kit (Thermo Scientific).

**Plasmid transformation.** To transfer the plasmid encoding for the desired protein, 40 µL BL21 gold *E. coli* cells (Novagen) (F-, ompT, hsdSB(rB- mB-), dcm+, Tetr, galλ (DE3) endA, Hte) were pipetted to 1 µL plasmid solution. The mixture was kept on ice for 15 min prior to a 20 sec heat-shock at 42°C. The cells were again kept on ice for about 5 min. After adding 400 µL super optimal medium with catabolite repression (SOC), the mixture was incubated for 1 h at 37°C. After the incubation, the cells were centrifuged at 800 x g for 3 min. 340 µL of the supernatant was discarded and the residual 100 µL of resuspended cells were plated onto LB agar plates (containing 100 µg/mL Amp and 1% glucose) using glass beads. The plates were incubated at 37°C over night (O/N).

**Protein expression.** To prepare a preculture, a single clone (in case of plasmid preparation) or 5 clones (in case of protein expression) was transferred to 20 mL of LB medium (containing 100 µg/mL Amp and 1% glucose) and incubated at 37°C O/N. The preculture was transferred into 1 L of TB medium (containing 100 µg/mL Amp) as main culture and incubated at 37°C until an optical density ($OD_{600}$) of 0.6 to 0.8 was reached. The main culture was cooled down and the expression was induced by adding 250 µL isopropyl β-d-1 thiogalactopyranoside (IPTG). The proteins were expressed at 20°C O/N. After about 16 h the main culture was transferred to centrifuge tubes from Beckmann Coulter and centrifuged in the Beckmann Coulter Avanti J20-XP with rotor JLA-8.100 at 4500 x g for 20 min. The supernatant was discarded, and the cells were resuspended in 20 mL His-wash buffer (200 mM KCl, 50 mM potassium phosphate, 30 mM imidazole, 10% glycerol, pH 7.0). To the suspension a small amount of Dnase I (deoxyribonuclease) (Sigma Aldrich) and 1 mM ethylenediaminetetraacetic acid (EDTA) was added. The cells were disrupted using the French Pressure Cell Press in one cycle. The obtained mixture was centrifuged using the JA 25.500 rotor at 40000 x g for 1 h. The supernatant was collected.

**Protein purification.** The crude extract was transferred to a His-NTA column (Takara) and washed with 5 CV (15 mL) of His-wash buffer. The protein was then eluted two times with 2.5 CV (7.5 mL) His-elution buffer (200 mM KCl, 50 mM potassium phosphate, 300 mM imidazole, 10% glycerol, pH 7.0) and transferred to a Strep-Tactin column. The column was washed with 2 CV (10 mL) Strep-wash buffer (250 mM potassium phosphate, 1 mM EDTA, 10% glycerol, pH 7.0). The protein was finally two times eluted with 3 CV (15 mL) Step-elution buffer (250 mM potassium phosphate, 1 mM EDTA, 10% glycerol, 2.5 mM desthiobiothin, pH 7.0). The obtained protein solution was concentrated and stored at -80°C until further purification. Proteins were further purified and analyzed after tandem affinity chromatography by SEC using the ÄKTA Basic system (GE Healthcare). The eluent was filtered, degassed and cooled before usage. The KS-MAT proteins were incubated for 1 h at 37°C prior to SEC to support dimerization. The protein sample was filtered using Ultrafree® Durapore® centrifugal filters (Merck). The KS-MAT constructs were purified using the Superdex 200 10/300 GL (cytiva) and Strep-wash buffer as eluent, whereas the ACP, MabA and Sfp proteins were purified using the HiLoad 16/600 Superdex 200 (cytiva) and ACP-SE buffer (50 mM potassium phosphate, 200 mM KCl, 10% glycerol, 1 mM EDTA, pH 7.0), MabA-SE buffer (50 mM sodium phosphate, 450 mM NaCl, 20% glycerol, pH 7.5) and Sfp-SE buffer (50 mM HEPES, 250 mM NaCl, 2 mM $MgCl_2$, 10% glycerol, pH 8.0) as eluent respectively. The column was equilibrated with 2 CV of buffer prior to usage. The protein sample was injected with automatic rinsing with two times the loop volume and eluted by 1.2 CV of buffer. All other settings were adjusted according to the manual.

**MabA Assay.** All solutions were prepared in 20 µL 384 well microplates (greiner bio-one) and measured using the CLARIOstar Plus platereader (BMG Labtech). The NADPH fluorescence was excited at 348-320 nm and detected at 476-420 nm. All solutions were prepared as 8 x stock in MabA buffer (50 mM sodium phosphate, 10% glycerol, pH 7.0), except of the Mal-X solution, which was prepared as 2 x stock. When adding the Mal-X solution (10 µL) the assay mixture was inverted thoroughly. The substrate titration experiment for wildtype characterization was performed with a knockout protein for a negative control at every condition. The reaction mixture without priming substrate was used as blanks for measurement of low activity (hydroxybutyryl-ACP and crotonyl-ACP) to take priming of decarboxylated malonyl-ACP into account. The fluorescence was measured during a period of 10 min. A final concentration of 5 µM MabA and 50 µM NADPH were used in all measurements. The final concentrations of enzyme and substrates in the assay depended on the experiment and is given in the figure descriptions.

**Acyl-ACP Synthesis.** To functionalize purified apo-ACP with acyl-moieties a Sfp phosphopantetheinylation was performed. 3 mM acyl-CoA, 600 µM apo-ACP and 30 µM Sfp in Sfp-reaction buffer (50 mM HEPES, 200 mM NaCl, 10 mM $MgCl_2$, pH 7.0) was incubated at 37°C for 20 min. (R)-Hydroxybutyryl-ACP was synthesized in respective solution with acetoacetyl-CoA (AA-CoA) and additional 30 µM MabA and 6 mM NADPH. To separate the product acyl-ACP from Sfp and acyl-ACP, Strep tactin affinity chromatography was performed. The solution was pipetted onto 5 mL of Strep-Tactin®XT. 4Flow® resin. The resin was subsequently washed with 3 CV Strep-wash buffer and acyl-ACP was eluted with 2.5 CV StrepXT-elution buffer (250 mM potassium phosphate, 1 mM EDTA, 10% glycerol, 50 mM biotin, pH 7.0). The elution fraction was rebuffered in ACP-buffer and analyzed with HPLC or urea PAGE. The analysis of different ACP species was carried out with HPLC on a Discovery®BIO Wide Pore C5 (Merck) with 0.1% TFA in water and 0.1% TFA in acetonitrile (39-42% $CH_3CN$ in 12 min with C2, C4 and crotonyl-ACP; 20-80% $CH_3CN$ in 15 min with C6 - C14-ACP). Around 0.25 nmol sample was injected to the column.

**Urea PAGE analysis.** Analysis of malonyl-ACP and hydroxybutyryl-ACP was performed with urea PAGE. Adapted from literature, the gel electrophoresis was carried out in a polyacrylamide gel containing urea.[51,52] The separating gel contained 15% acrylaminde/ bisacrylamide, 1.12 M Tris, 7.5 M urea, 0.1% (v/v) N,N,N',N'-tetramethylethyldiamine and 0.03% (w/v) ammonium persulfate. The loading dye (2x) contained 1% bromphenol blue, 25% glycerol, 62.5 mM Tris, 2 M urea and either 10 mM N-ethylmaleimide (NEM) or N-aminoethylmaleimide (NAM). Samples were incubated for 10 min in loading dye prior to loading to the gel. The electrophoresis was performed at 70 V for 15 min and subsequent 200 V for 1 – 1.5 h with urea-page running buffer (25 mM Tris, 200mM glycine), cooling was applied. The gel was stained in an aqueous solution of 0.1% Coomassie Blue R-250, 50% methanol and 10% acetic acid over-night. The background was destained with an aqueous solution of 10% ethanol and 10% acetic acid.

For the qualitative analysis of the KS catalyzed reaction with urea PAGE, 200 nM KS, 50 µM Mal-ACP and 50 µM C10-ACP or 1.5 µM KS, 180 µM Mal-ACP and 50 µM HB-ACP in MabA buffer were incubated at 25°C. The reaction was stopped by adding equal volume of isopropanol to precipitate the KS. After centrifugation at 20000 x g for 10 min, the supernatant was analyzed as described above.

**Hexanyol-SNAC synthesis.** This synthesis was adopted from Peter et al. and Valenzano et al.[53,54] 0.6 mL triethylamine (4.3 mmol, 2 eq.) was added to the hexanoic acid (4.3 mmol, 2 eq.) in 20 mL tetrahydrofurane (THF) (dry) at 0°C. 0.4 mL ethyl chloroformate (4.3 mmol, 2 eq.) was added and the suspension was stirred excessively under argon for 45 min at 0°C. The suspension was filled into 50 mL tubes and centrifuged for 10 min at 4°C and 3220 x g to remove insoluble salts. About 20 mL of the supernatant was added to 230 mL SNAC (2.2 mmol, 1 eq.) in 10 mL 0.1 M NaHCO3 (pH 8) and stirred for 1 h at room temperature. The product was extracted 3-4 times from the resulting solution with 30 mL diethyl ether each time. The organic phase was washed 3 times with 10 mL 0.1 M $Na_2CO_3$ (pH 9) each time and 5 mL sodium chloride solution (sat.), dried with magnesium sulfate and filtered. The solvent was removed using a rotary evaporator. The raw product was further purified by column chromatography using EE:H 1:1 to 3:1. Hexanoyl-SNAC was obtained as colorless solid (89 mg, 20%, $R^f$ (EE:H 4:1) = 0.26, m/z [M+Na] = 240.04). 250 MHz, $CDCl_3$ $\delta$[ppm]: 5.68 (s, NH), 3.44 (td, $^3J_{HH}$=6.1 Hz, 2H, $CH_2$-2), 3.02 (t, $^3J_{HH}$=6.4 Hz, 2H, $CH_2$-3), 2.57 (t, $^3J$ HH=7.5 Hz, 2H, $CH_2$-4), 1.97 (s, 3H, CH3-1), 1.66 (quint, $^3J_{HH}$=7.3 Hz, 2H, $CH_2$-5), 1.36 - 1.25 (m, 4H, $CH_2$-6 - $CH_2$-7), 0.90 (t, $^2J_{HH}$=6.8 Hz, 3H, $CH_3$-8). A contaminant between 1.6 and 1.8 ppm could not be identified. The interpretation of measured NMR spectra were supported by the literature.[55,56]

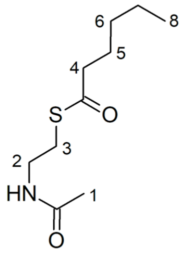


**Acyl-CoA synthesis.** This synthesis was adopted from Peter et al. and Valenzano et al.[53,54] The respective acid (0.31 mmol, 6 eq.) was dissolved in 2 mL THF and cooled to 0°C. 40 µL triethylamine (0.29 mmol, 6 eq.) and 27 µL chloroformate (0.31 mmol, 6 eq.) were pipetted to the solution and stirred for 45 min at 0°C under argon. The reaction mixture was transferred to a 2 mL microcentrifuge-tube and centrifuged for 5 min at 20000 x g to get rid of insoluble salts. The supernatant was transferred to 40 mg CoA (49 µmol, 1 eq) in 2 mL 0.1 M sodium hydrogen carbonate solution. The mixture was stirred for 1 h at room temperature. The reaction mixture was poured into 25 mL of cold acetone (-20°C) and centrifuged for 5 min at 10000 x g. The raw products were obtained as colorless solids and stored at -20 °C until further purification. To isolate the CoA-esters, the samples were dissolved in 200 mM ammonium acetate at pH 6 and centrifuged for 2 min at 20000 x g to separate any solids. The product solution was injected to the Acclaim. Polar Advantage II LC-Column (Thermo Scientific) and eluted with a buffer (200 mM ammonium acetate, pH 6)-acetonitrile gradient (10 - 60% acetonitrile). The combined product fractions were concentrated under vacuum. The product was obtained at a solution in buffer. The yield was determined by measuring the concentration of the product solution. The products were analyzed using ESI-MS and HPLC. Acetyl-CoA: 13 nmol, 26%, m/z $[M+H]^+$ = 810.40 / Butyryl-CoA: 13 nmol, 26%, m/z $[M+H]^+$ = 838.08 / Hexanoyl-CoA: 16 nmol, 33%, m/z $[M+H]^+$ = 866.20 / Octanoyl: 18 nmol, 36%, m/z $[M+H]^+$ = 894.19 / Decanoyl-CoA: 15 nmol, 31%, m/z $[M+H]^+$ = 922.22.

**Thermal fluorescence assay.** The stability of mutants was assayed in five different buffers that are used throughout the purification and evaluation procedure. A solution of around 2 µg protein in His-wash buffer, His-elution buffer, Strep-wash buffer, strep-elution buffer and MabA buffer was prepared on ice in 96-well plates (Multiplate® PCR Plates™, Bio-Rad) and Sypro™ Orange (Invitrogen) was added according to manufacturer's guideline. The well plate was sealed with optical tape (iCycler iQ®, Bio-Rad) and centrifuged shortly at 3000 xg. The measurements were performed in a real-time thermocycler (CFX Connect™, Bio-Rad) at excitation/emission wavelengths of 450-490/650-580. A stepwise temperature increase of 0.5°C/30s was performed between 5°C and 95°C.

**System Preparation.** The crystallographic x-ray structure of murine FAS KS-MAT domain (PDB ID: 6ROP, 2.70 Å resolution)[24] was used to prepare the KS-substrate models used in this study. The asymmetric unit comprises four copies of the KS-MAT di-domain arranged in two functional dimers MAT-KS:MAT-KS (chains A-B:C-D).
Two adjacent KS domains (residues 1-409 plus 825-852) from chains A and B were extracted and used to build the studied model. There were no water molecules present in the structure. Furthermore, this structure had an octanoyl moiety covalently bounded to the KS domain that was deleted before modeling the substrate in the KS active site.
The acetyl moiety carbonyl oxygen was oriented to the backbone amides of Cys161 and Phe395, which are expected to act as an oxyanion hole during the KS-catalyzed reaction.[22,24] The catalytic cysteine (Cys161) was bonded to the octanoyl moiety in the x-ray structure. This residue was oriented to His331 to approximate the KS active site to a catalytically competent conformation in the prepared system.
The H++ webserver[57] was used to predict the protonation states of the titratable residues. According to H++, Cys161 should exist in neutral form (-SH). The protonation of the histidines was carefully manually checked, and the predictions from H++ were accepted except for His293, which was protonated in its δ nitrogen.
The coordinates from PaM10 were saved and transferred to the 6ROP active site. The GaussView software was used to convert the PaM10 into a Ppant group and to add an acetyl moiety to its terminal thiol group, generating the acetyl-Ppant substrate. The coordinates of the introduced acetyl moiety were designed to maximize the hydrogen bonds between its carbonyl oxygen and the backbone amides of Cys161 and Phe395, which are expected to form an oxyanion hole throughout the KS-catalyzed reaction. It is worth noting that substrates containing butyryl, C6acyl, C10acyl, crotonyl, and hydroxybutyryl were also created using the same methodology to obtain Ppant substrates.
The enzyme residues were parameterized using the AMBER ff14SB force field. The substrates were parameterized using the following protocol: First, they were divided into two distinct molecules, Ppant and the substrate moieties, which were parameterized as independent units. Hydrogen atoms were added to the carbonyl carbon of the substrate molecules and to the terminal sulfur atoms of the Ppant group to complete their valence shells. The Antechamber module of the Amber 18 package was used to parameterize the independent units with GAFF2. The charges were derived from a RESP fitting of the electrostatic potential determined at HF/6-31G(d) level of theory. The charge of the added hydrogen atoms was kept equal to zero and constant throughout the calculation of the electrostatic parameters for each molecule. The Xleap module of the Amber 18 package was used to delete the added hydrogen atom and to create a covalent bond between the substrate moiety and Ppant molecules. Xleap was also used to add $Na^+$ counterions and solvate the system with an octahedral box of TIP3P water molecules within a radius of 12 Å from the surface of the protein.

**MD Simulations for substrate binding.** The energy of the prepared systems was minimized to alleviate steric clashes or unfavorable tensions that may be present in the modeled systems. Gromacs 2021 was employed, and the minimization was performed in two steps. First, the water molecules, hydrogens, and counter-ions were minimized using the steepest descent algorithm. Periodic boundary conditions were imposed to account for long-range interactions. A radius of 10 Å was defined as the cut-off distance for short-range electrostatic and Lennard-Jones interactions. The entire system was minimized in the following stage using the same conditions.
The system was then equilibrated, starting with a 100 ps simulation with an NVT ensemble using the modified Berendsen thermostat and a reference temperature of 300 K. Then, a 100

ps equilibration MD was run with the NPT ensemble, in which the density of the system was equilibrated at 300 K and 1 bar using the modified Berendsen thermostat and the Berendsen barostat. During these stages, the protein and the substrates were constrained with positional restraints of 1000 kJ $mol^{-1}$ $nm^{-2}$ and 2000 kJ $mol^{-1}$ $nm^{-2}$, respectively.
The equilibration phase was followed by a 50 ns simulation run with the NPT ensemble in which the hydrogen atom H17 and the oxygen atom O7 from Ppant and three Thr residues (Thr262, Thr295, and Thr297) from the protein active site were restrained with a force of 1000 kJ $mol^{-1}$ $nm^{-2}$, to guarantee the initial orientation of the substrate through the establishment of hydrogen bonds between the threonines and the atoms from Ppant. Finally, an unrestrained production phase of 50 ns using the NPT ensemble was conducted.

**Modelling of the Rattus norvegicus FAS.** The Rattus norvegicus FAS (rFAS) that served as the foundational template was sourced from AlphaFold (AF) with the corresponding code AF-P12785-F1. As the AlphaFold entry revealed the specific residues responsible for forming the linker between the Ketoacyl Reductase (KR) and Acyl Carrier Protein (ACP) domains, homology modelling was implemented to construct the linker.
First, the thioesterase (TE) and ACP domains were deliberately excluded from the AF-P12785-F1 template. The ACP domain was subsequently incorporated back into the structure using an AlphaFold model of the KS:ACP complex, obtained from an AlphaFold Colab calculation.[58] Except for the absent TE domain, the target FAS sequence was submitted to the SWISS-Model software for model construction.[59]
The geometry of the modelled FAS was then optimized with the Sander module of the Amber software package through 1000 steepest descent steps and 1500 conjugate gradient steps.[60]

**Parameterization of the Rattus norvegicus FAS.** The final system was built on the tleap program using an AMBER ff14SB force field and TIP3P water molecules. A cuboid box of water molecules with faces at a minimum distance of 10 Å from the protein was created to solvate the protein. Furthermore, 61 Na+ counterions were added to neutralize the system's total charge. Given the large size of the modelled FAS protein (4404 residues) and the simulation box necessary to simulate it, the total system was very large, consisting of 924340 atoms, limiting the length of the subsequent MD simulations.
The protonation state of ionizable residues was estimated using the web server PDB2PQR to predict their local pKa values. The AMBER parameter files were converted to GROMACS parameters using a Python script and the amber.python command, as the molecular dynamics simulation was conducted using the GROMACS software.

**MS Simulations for ACP binding.** A molecular dynamics simulation of 200 ns was conducted using the GROMACS software. At first, the system underwent a two-stage minimization process, initially addressing the water molecules and then minimizing the entire system. Subsequentially, the complex was equilibrated for 20 ns in the canonical (NVT) ensemble with restraints applied to all the protein atoms except the ACP:KR linker. In the next step, an equilibration was performed with positional restraints imposed on the ACP:KS interfacial residues (45-49, 198-205, 297-298 of KS and 47-74 of ACP) lasting 200 ns. In both equilibration phases, conducted within the NVT ensemble, a V-rescale (modified Berendsen) thermostat was utilized. The subsequent and final molecular dynamics run was done at the isothermal-isobaric (NPT) ensemble, using the Berendsen barostat without positional restraints, for 200 ns. Therefore, a total simulation time of 420 ns was run.
The reference temperature for all MD simulations was set to 310.15 K.
In one of the replicates, the NPT run was prolonged up to 1 µs employing the Parrinello-Rahman thermostat.
Therefore, a total of 1.84 µs of MD simulation was run. RMSD values were then calculated using analysis tools within the GROMACS software.

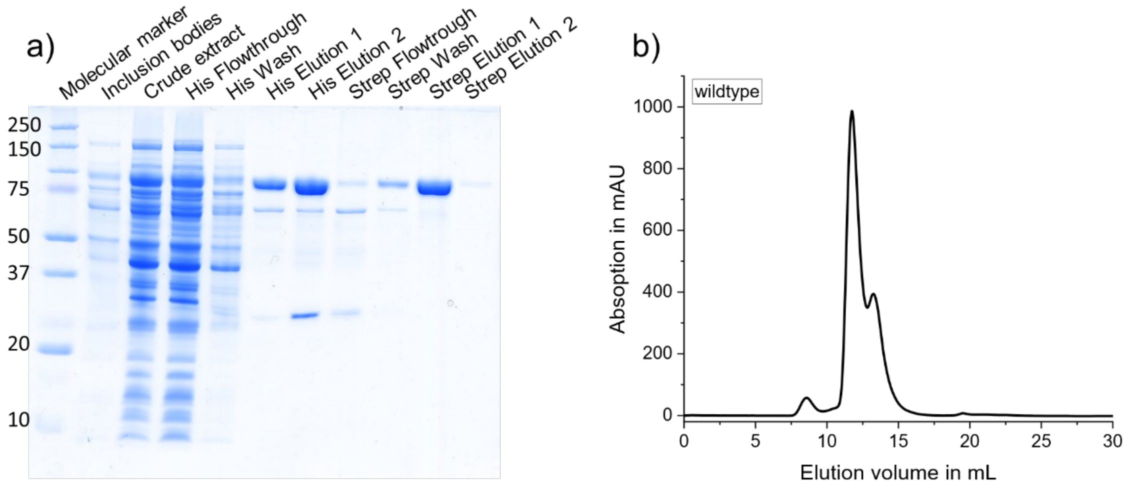


***S1 Purification and quality control of KS-MAT$^{S581A}$.*** *a) SDS PAGE of of the purification procedure using tandem affinity chromatography. The target protein has a size of around 97 kDa. b) Size exclusion chromatogram shows aggregation, that elute at around 8 mL, dimer at around 11 mL and monomeric KS-MAT at around 13 mL.*

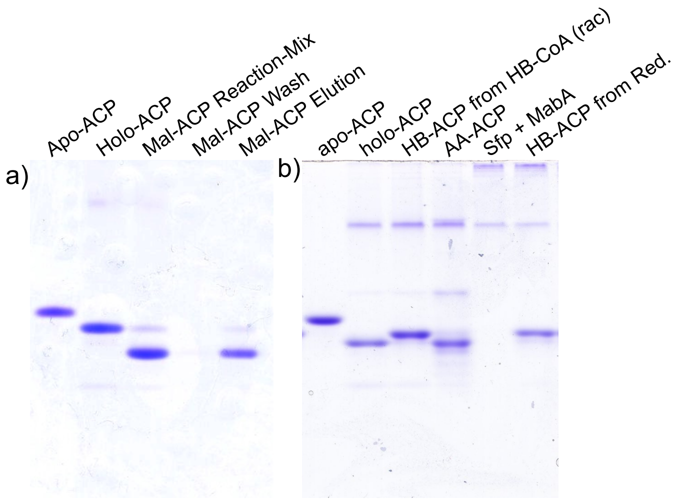


***S2 Urea polyacrylamide gels of X-ACPs.*** *a) NEM was included in the loading dye. Apo-ACP and holo-ACP are shown as reference. The reaction-mix of Mal-ACP generation includes apo-ACP, Sfp and Mal-CoA. The wash fraction and the elution fraction according to the purification protocol are shown. Mal-ACP was acquired in 90% purity with small amounts of holo-ACP from thioester hydrolysis. b) NAM was included in the loading dye. Apo-ACP and holo-ACP are shown as reference. HB-ACP was generated in a Sfp catalyzed reaction from apo-ACP and HB-CoA (rac). AA-ACP was generated from apo-ACP and AA-CoA. Finally, (R)-HB-ACP was generated by phosphopantetheinylierung from apo-ACP and AA-CoA with subsequent reduction of MabA with NADPH. (R)-HB-ACP was acquired in around 90% purity.*

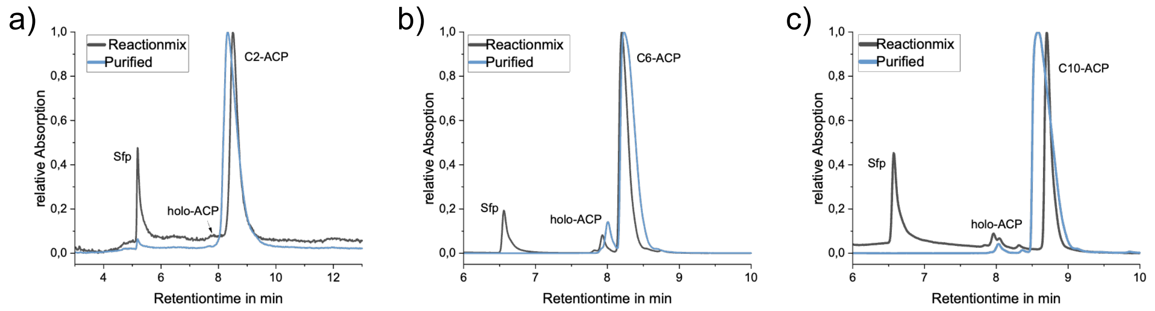


***S3 HPLC chromatograms of Acyl-ACPs**. a) Acetyl-ACP b) Hexanoyl-ACP and c) Decanoyl-ACP. Black: Reaction mix of acyl-ACP generation including Sfp, apo-ACP and respective acyl-CoA. Blue: Purified acyl-ACP after strep-tactin-column purification and rebuffering in ACP buffer. Peaks of acyl-ACP are normalized.*

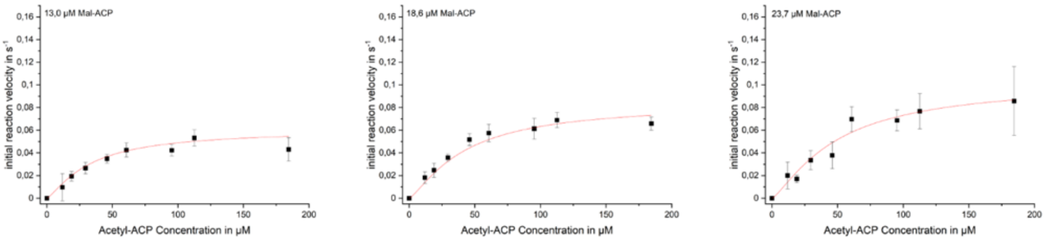


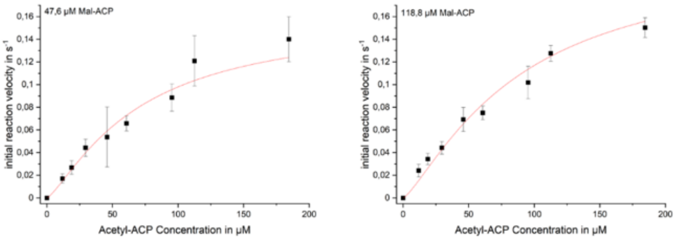


| Constant | Acetyl-ACP | Malonyl-ACP |
|---|---|---|
| $K_m$/ µM | 100 ± 26 | 33 ± 7 |
| $K_{cat}$/ $s^{-1}$ | 0,26 ± 0,05 | 0,26 ± 0,05 |
| $K_{cat}/K_m$ | 0,0026 ± 0,0008 | 0,007 ± 0,002 |
| h | 1,28 ± 0,11 | 1,28 ± 0,11 |

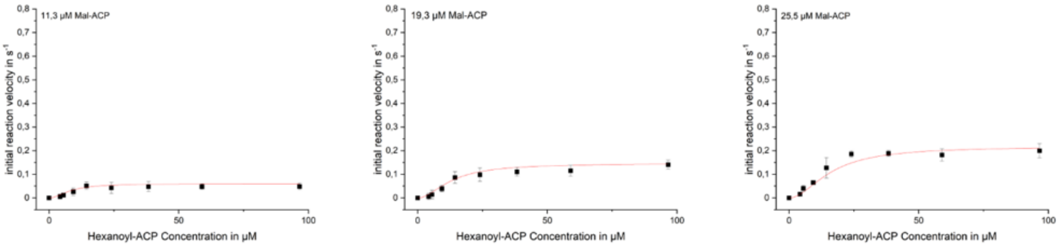


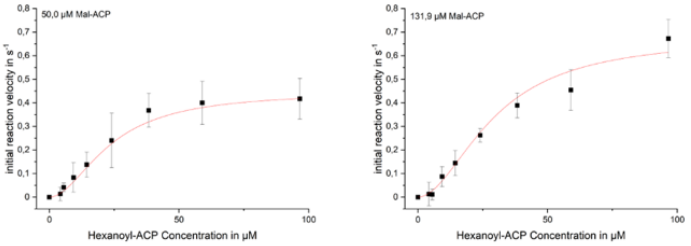


| Constant | Hexanoyl-ACP | Malonyl-ACP |
|---|---|---|
| $K_m$/ µM | 31 ± 3 | 41 ± 3 |
| $K_{cat}$/ $s^{-1}$ | 0,75 ± 0,05 | 0,75 ± 0,05 |
| $K_{cat}/K_m$ | 0,024 ± 0,003 | 0,018 ± 0,002 |
| h | 1,90 ± 0,11 | 1,90 ± 0,11 |

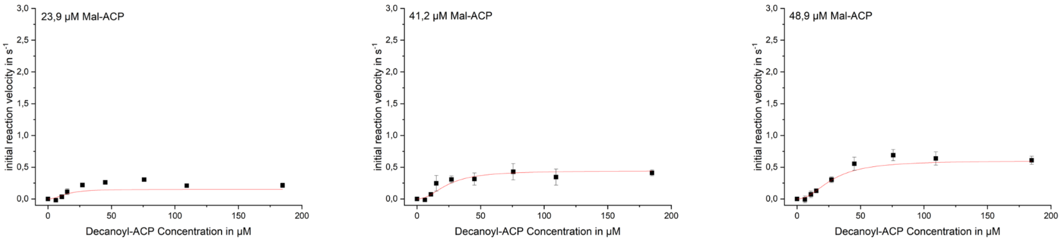


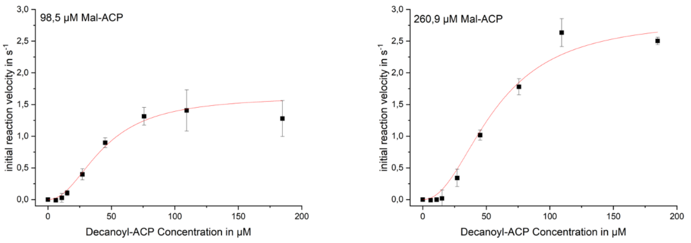


| Constant | Decanoyl-ACP | Malonyl-ACP |
|---|---|---|
| $K_m$/ µM | 60 ± 5 | 97 ± 8 |
| $K_{cat}$/ $s^{-1}$ | 3,2 ± 0,2 | 3,2 ± 0,2 |
| $K_{cat}/K_m$ | 0,053 ± 0,006 | 0,033 ± 0,003 |
| h | 2,1 ± 0,1 | 2,1 ± 0,1 |

***Figure S4: Kinetic characterization of the wildtype KS using the MabA assay.*** *The initial reaction velocity is plotted against eight different concentrations of acyl-ACP at five malonyl-ACP concentrations. The KS concentration for all measurements was 300 nM. Individual measurement series were performed for each of three carbon chain lengths. The obtained titration curves were globally fitted for each chain length using the Hill equation without any constraints. The kinetic constants are provided in the respective tables. The enzymatic efficiencies* $k_{cat}/K'$ *are given in* $s^{-1}\mu M^{-1}$*. a) acetyl-ACP b) hexanoyl-ACP c) decanoyl-ACP*

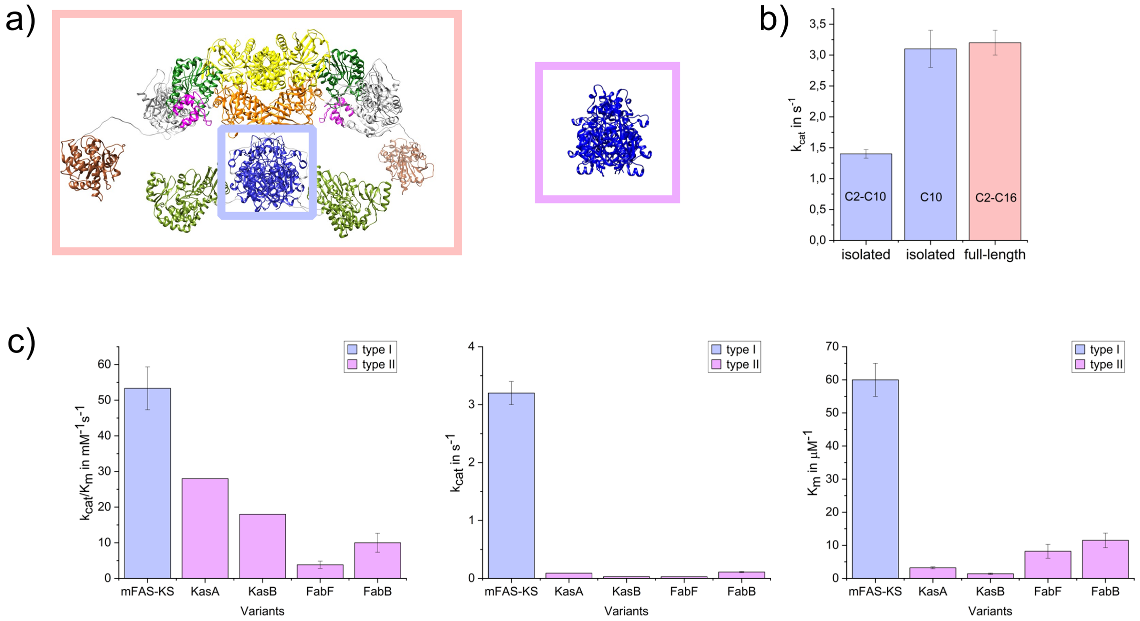


***Figure S5 Enzymatic constant comparison*** *a) Schematic depiction of mFAS (orange), mFAS-KS (blue) and type II KS (pink) (PDB: 1G5X)*[23,61] *b) comparison of $k_{cat}$ of isolated mFAS-KS (blue) and full-length FAS (orange). c)-e) Enzymatic constants of isolated mFAS-KS (blue) and type II bacterial KS (pink). Apparent enzymatic constants of KasA and KasB refer to C16-ACP, FabF and FabB data were obtained with C14-ACP.* [35,45]

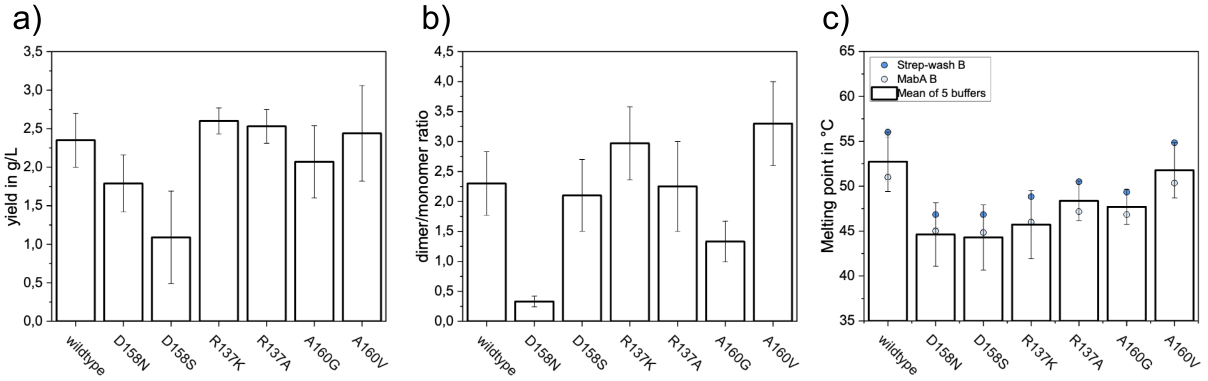

d)

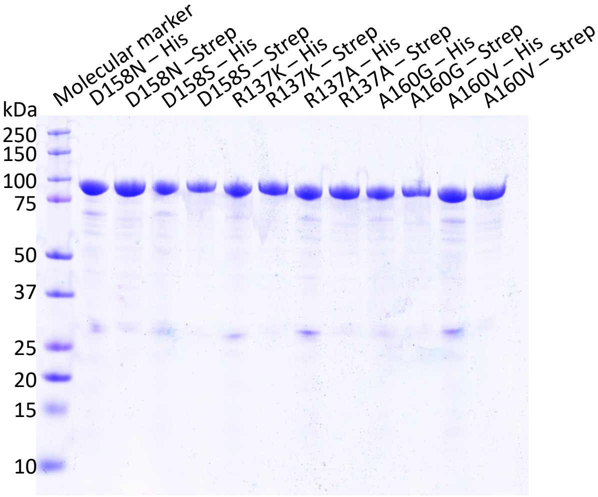


e)

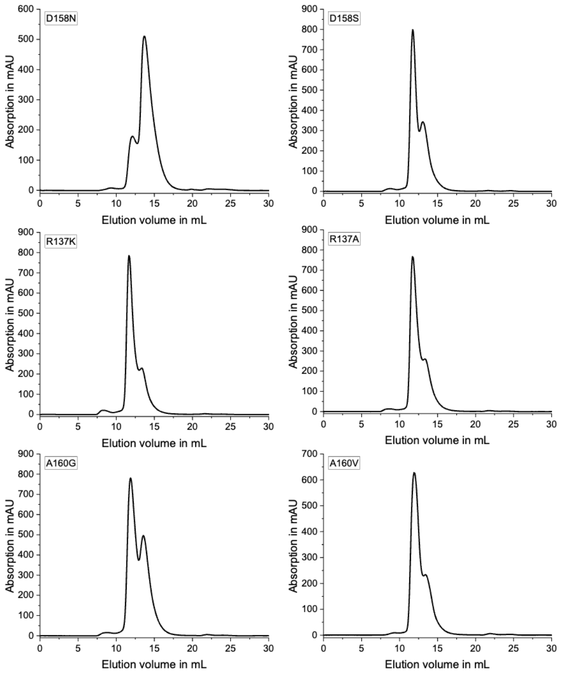


***Figure S6: Analysis of the protein quality of mutants that were used to investigate the enzyme allostery**. a) The average of the protein yields of three expressions in mg protein per liter of main culture. b) The average of dimer/monomer ratio obtained for three biological replicates from the size exclusion chromatogram. c) Thermal fluorescence assay was performed for all mutants in five buffers, which are used in the workflow of protein preparation and kinetic evaluation. The minimum of the melting point derivative is given as melting point. The storage buffer (Strep-wash B) and assay buffer (MabA B) are shown as separate data points. All datapoints represent biological triplicates. d) Exemplary SDS PAGE analysis of the elution fractions of of Ni-NTA column and Strep-Tactin column of all mutants. e) Representative size-exclusion chromatograms of all mutants.*

*Table S6: The values derived from the titration experiment of mutants, that attempted to delete the hydrogen bond network and thus intersubunit communication.*

| Protein | h | $V_{max}^{app.}$ / $s^{-1}$ | $K_m^{app.}$/ µM |
|---|---|---|---|
| wildtype | 2.9 ± 0.9 | 1.2 ± 0.1 | 27 ± 3 |
| D158N | not active | | |
| D158S | not active | | |
| R137K | 3.2 ± 0.7 | 1.4 ± 0.1 | 32 ± 3 |
| R137A | 3.0 ± 0.9 | 0.38 ± 0.04 | 26 ± 3 |
| A160G | 1.8 ± 0.6 | 0.6 ± 0.3 | 60 ± 31 |
| A160V | not active | | |

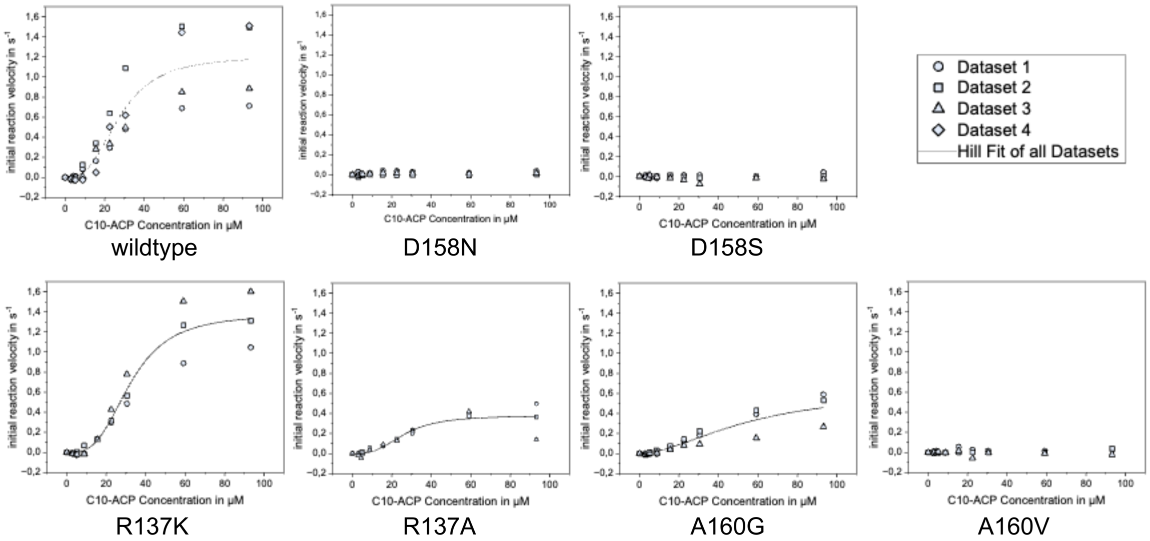


***S7 Titration of C10-ACP in titration measurements with mutants**. The measurements were performed with 300 nM KS at 50 µM Mal-ACP. The data was fitted with the Hill equation using OriginLab®. The results of the fit can be found in table S6.*

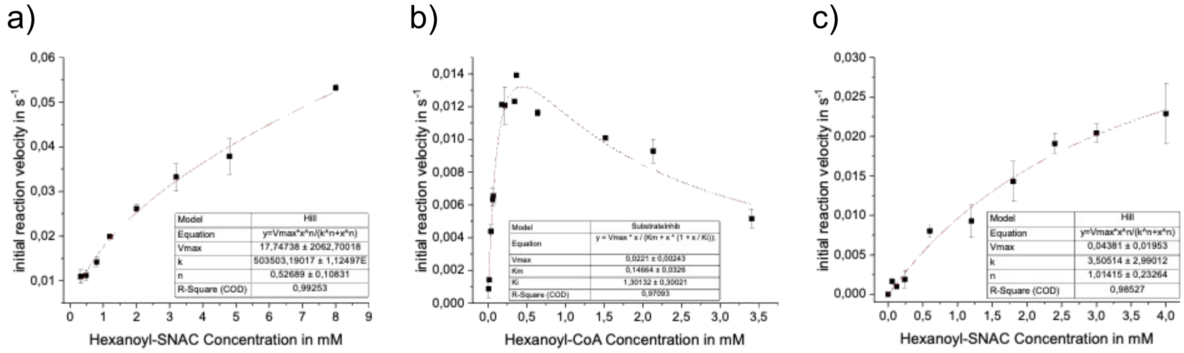


***S8 Carrier Analysis** a) The wildtype KS was assayed at 1.7 µM protein and 250 µM malonyl-CoA. The data was to the Hill equation to take a potential cooperativity into account. b) The wildtype KS was assayed at 1.9 µM protein and 470 µM malonyl-CoA. The data was fitted to the substrate inhibition equation, because auf significant activity decrease with increasing substrate concentration. c) The wildtype KS was assayed at 1.4 µM protein and 47 µM malonyl-ACP. The data was fitted to the Hill equation to take potential cooperativity into account. All data fits were performed using OriginLab®.*

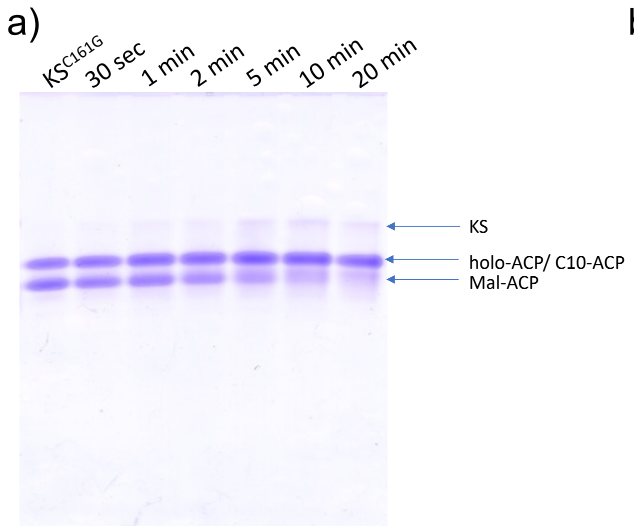


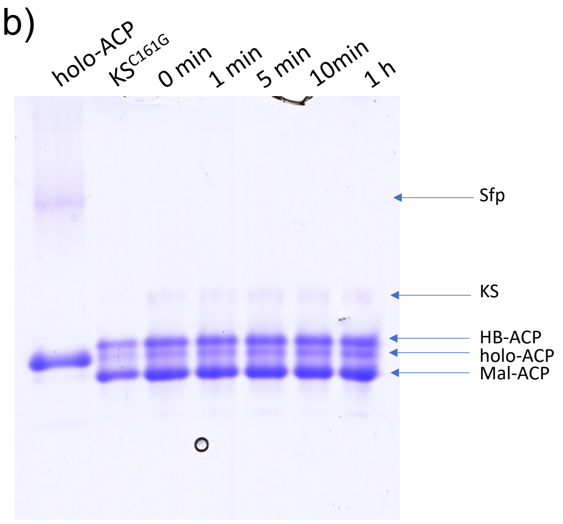


***S9 Urea PAGE analysis of KS catalyzed condensation reaction.*** *a) The gel shows the time course of the reaction of decanoyl-ACP with malonyl-ACP in presence of KS. Negative control includes KS$^{C161G}$ as a functional knockout. Note that decanoyl and holo-ACP cannot be separated under the chosen conditions. The reaction progress is monitored by the amount of malonyl-ACP. b) The gel shows the time course of the reaction of (R)-hydroxybutyryl-ACP with malonyl-ACP in presence of KS. Holo-ACP is shown as reference. No depletion of HB-ACP nor malonyl-ACP can be seen.*

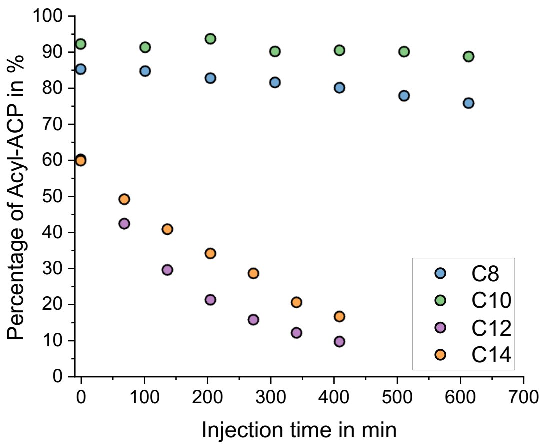


***S10 Hydrolysis of Acyl-ACPs***. *Freshly prepared Acyl-ACPs were analyzed after different time with HPLC. The peak integrals of octanoyl-ACP, decanoyl-ACP, lauryl-ACP and myristyl-ACP are shown in dependance of the hydrolysis time. The samples were kept tempered at 4°C during this measurement series.*

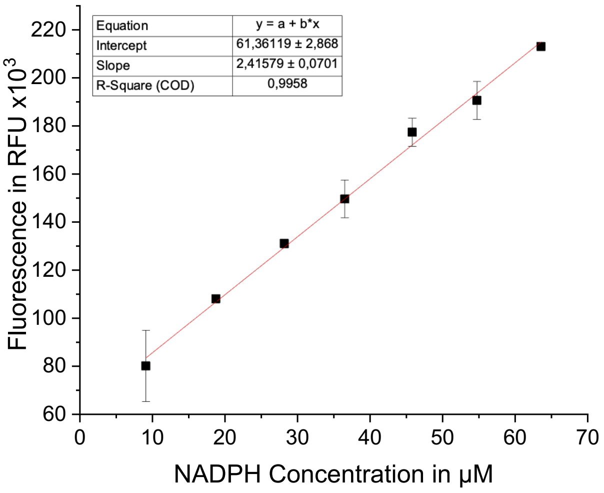


***S11 Representative calibration***. *Fluorescence of NADPH in dependance of NADPH concentration in measured with the same settings and under the same conditions as the final KS activity measurements.*

***Table S1. The linker length between the ACP and KR domains for both rFAS chains***. *The distance between the residue's alpha-carbon at the start and end of the linker was computed. Mean values and standard deviations are provided for each chain. The differences in chain length between the replicates illustrates the conformational flexibility of the rFAS multienzyme and the diversity of structural solutions for binding both ACP domains to KS simultaneously.*

| Replica | Chain A (Å) | Chain B (Å) |
|---|---|---|
| 1st Replica | 43.1 ± 0.9 | 45.5 ± 0.8 |
| 2nd Replica | 38.3 ± 0.8 | 33.2 ± 1.0 |
| 3rd Replica | 34.5 ± 0.6 | 30.0 ± 0.6 |

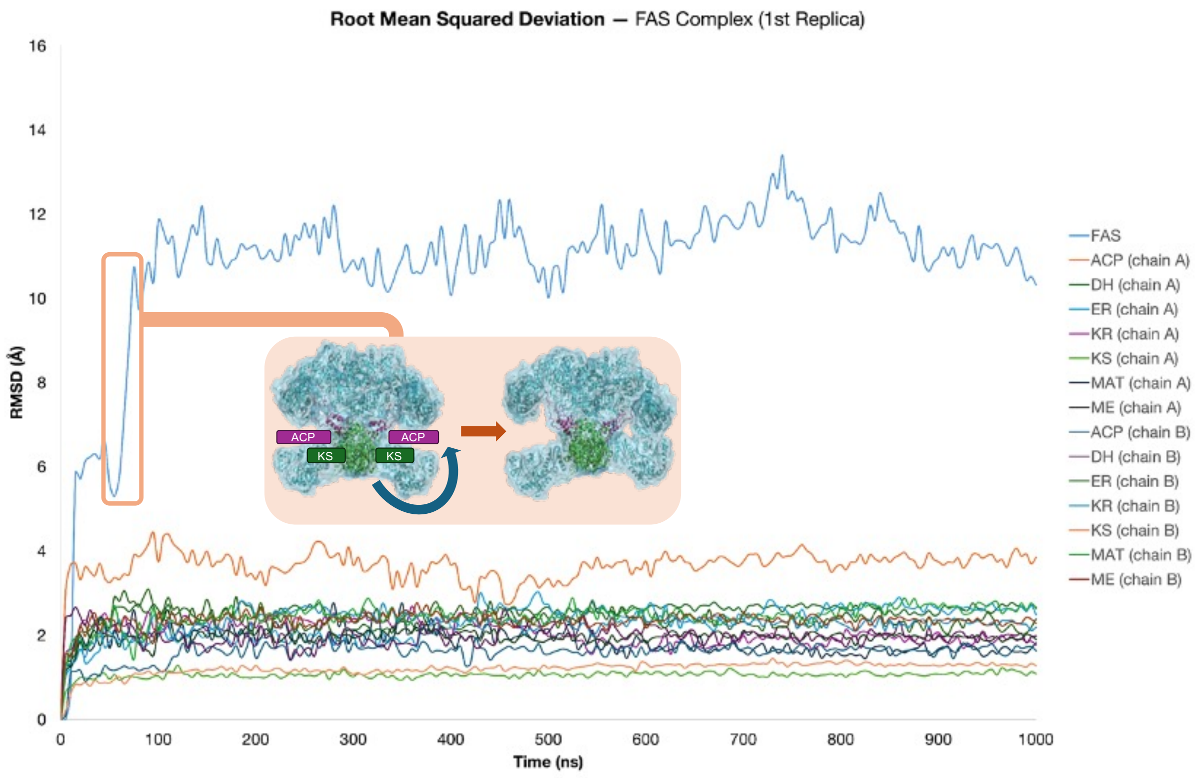


***Figure S12.*** *The Root Mean Square Deviation (RMSD) of the FAS complex and its individual domains from both polypeptide chains in the first replica. Running averages of the RMSD values were calculated to facilitate the visual analysis of the complex and its domains' stability. The different simulation steps are depicted, emphasizing the conformational shifts observed during the equilibration simulations. The graphic shows that the rFAS adopts an asymmetrical conformation, simultaneously binding both ACP domains to KS. The asymmetric conformation is reached through rigid-body domain motions, as confirmed by the low RMSD of the individual domains. A visual representation of the conformational shift is provided for clarity, with the ACP and KS domains highlighted in magenta and green, respectively.*

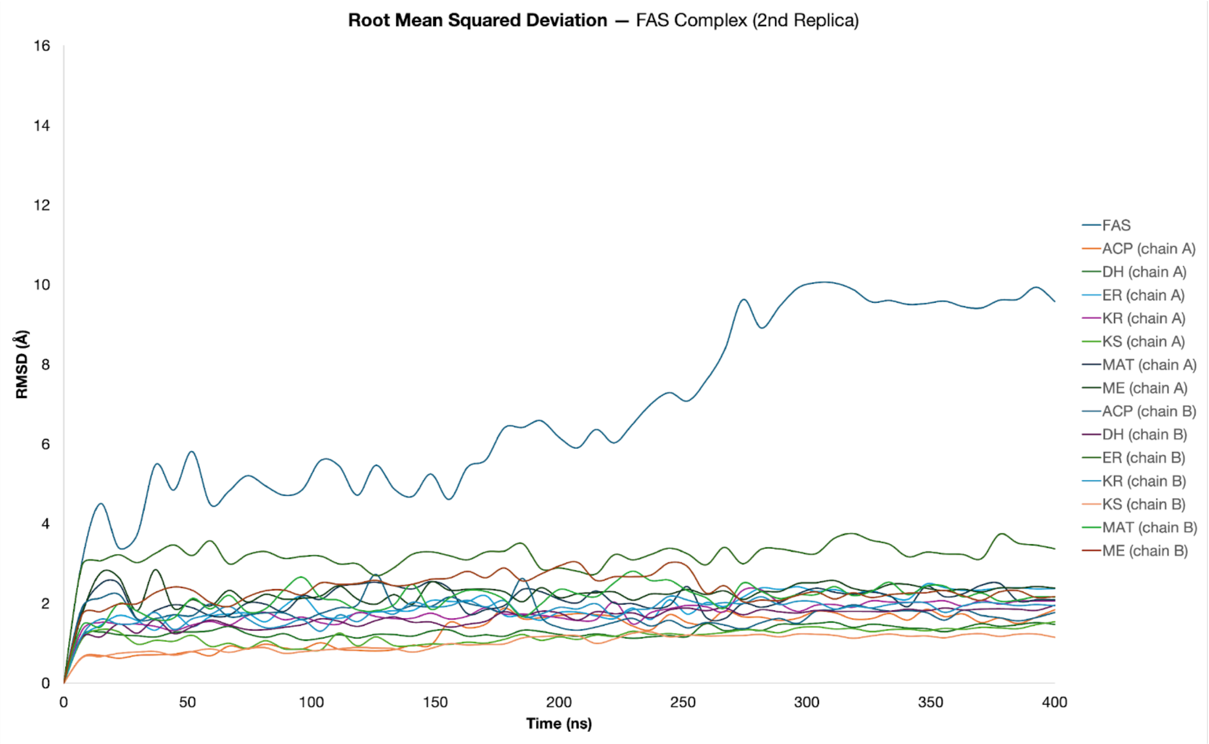


***Figure S13***. *The Root Mean Square Deviation (RMSD) of the FAS complex and its domains from both polypeptide chains in the second replica. Running averages of the RMSD values were calculated to facilitate the visual analysis of the complex's and domains' stability.*

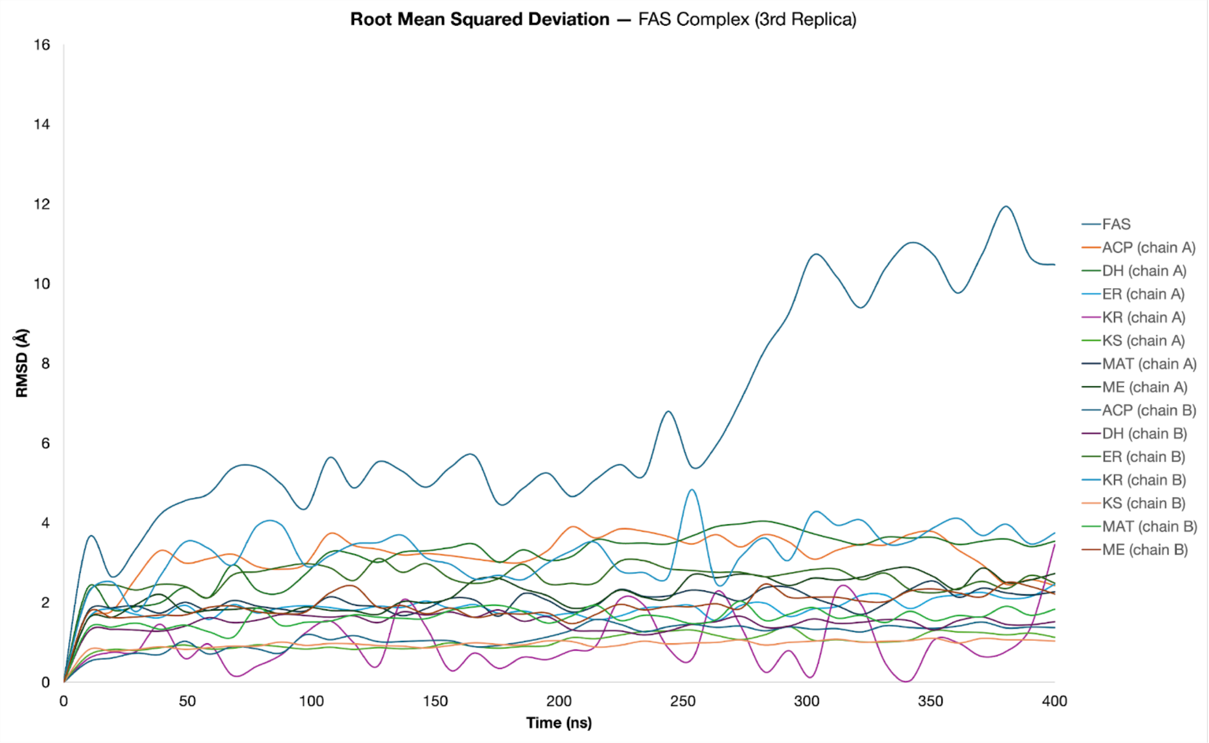


***Figure S14*** *The Root Mean Square Deviation (RMSD) of the FAS complex and its domains from both polypeptide chains in the third replica. Running averages of the RMSD values were calculated to facilitate the visual analysis of the complex and its domain stabilty. In this third replica, as ACP dissociates from KS, the RMSD values exhibit reduced stability, indicating increased flexibility of ACP within the overall system. The dissociation observed after the conformational shift in the third replica is within expectation, as ACP is anticipated to dissociate easily to facilitate the continuation of the FAS cycle.*

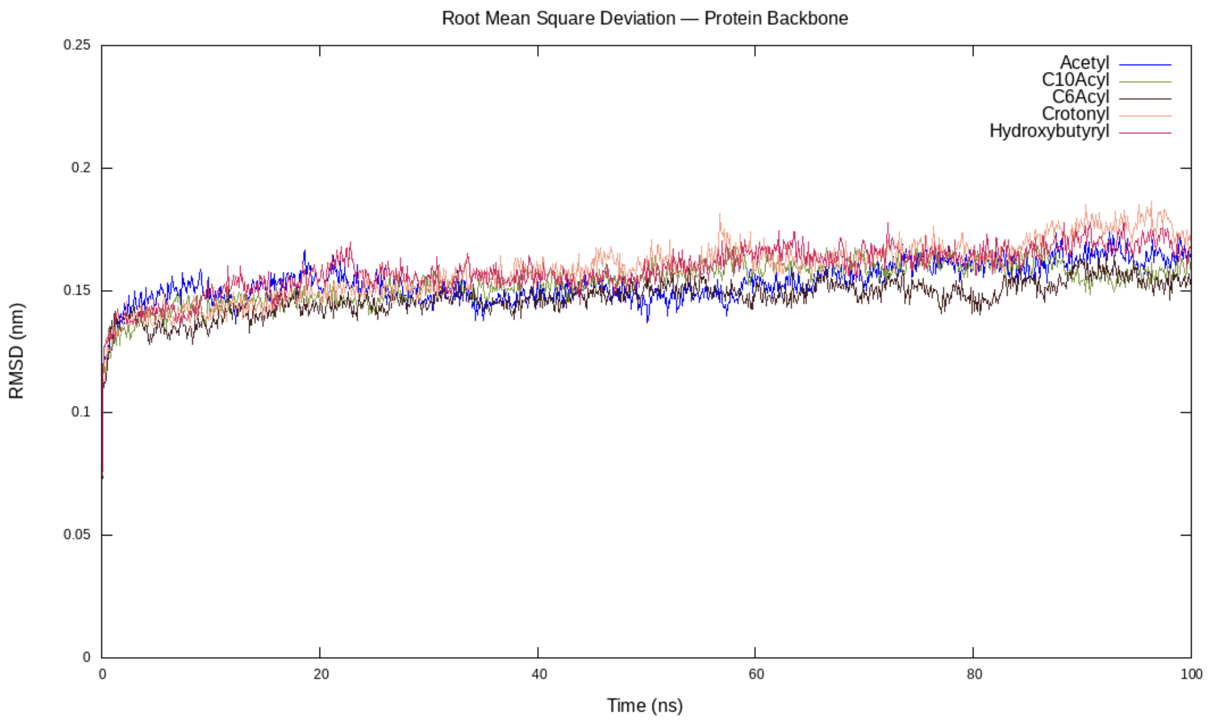


***Figure S15 The Root Mean Square Deviation (RMSD) of the KS protein backbone in each KS:substrate complex throughout the 100 ns simulations.*** *The KS:acetyl complex is colored in blue, KS:C10acyl complex in green, KS:C6acyl complex in brown, KS:crotonyl complex in pink and KS:hydroxybutyryl in magenta. RMSD values are represented in nm. The positional restraints that assured the substrate initial position were released in the 50 ns mark.*

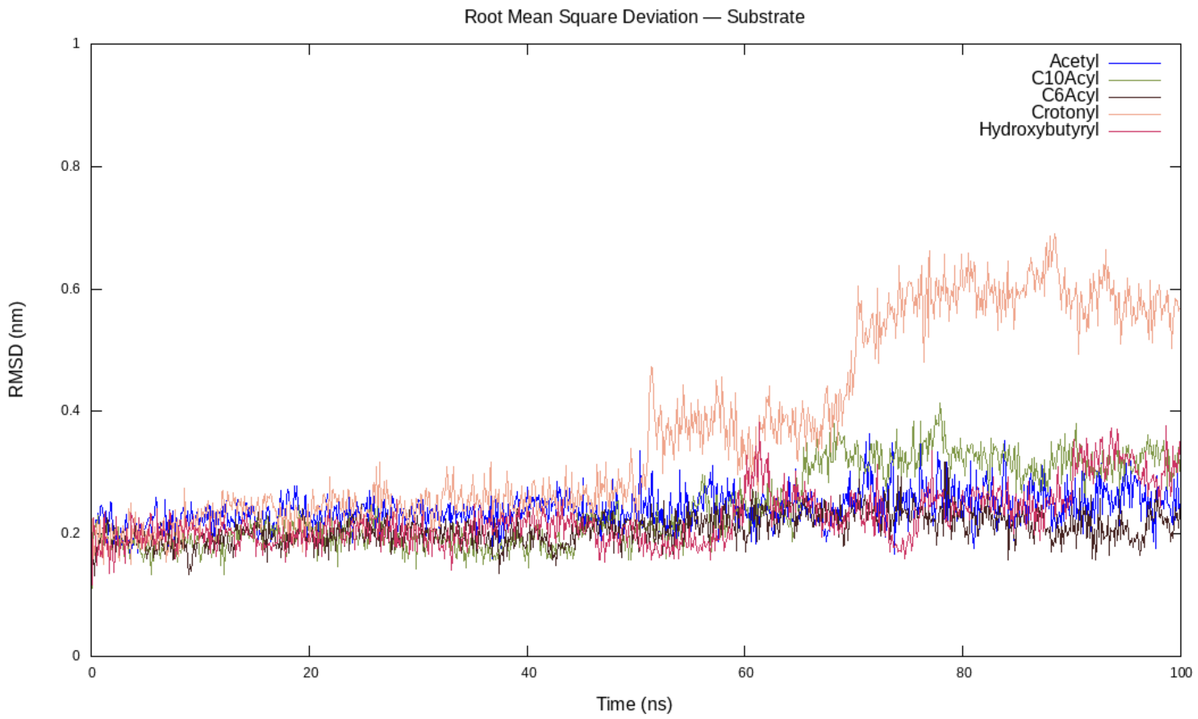


***Figure S16 Root Mean Square Deviation (RMSD) of each substrate in every KS:substrate complex throughout the 100 ns simulation.*** *The KS:acetyl complex is colored in blue, the KS:C10acyl complex in green, the KS:C6acyl complex in brown, the KS:crotonyl complex in pink, and the KS:hydroxybutyryl in magenta. RMSD values are represented in nanometers. It is worth noting that the positional restraints that assured the substrate initial position were released in the 50 ns mark.*

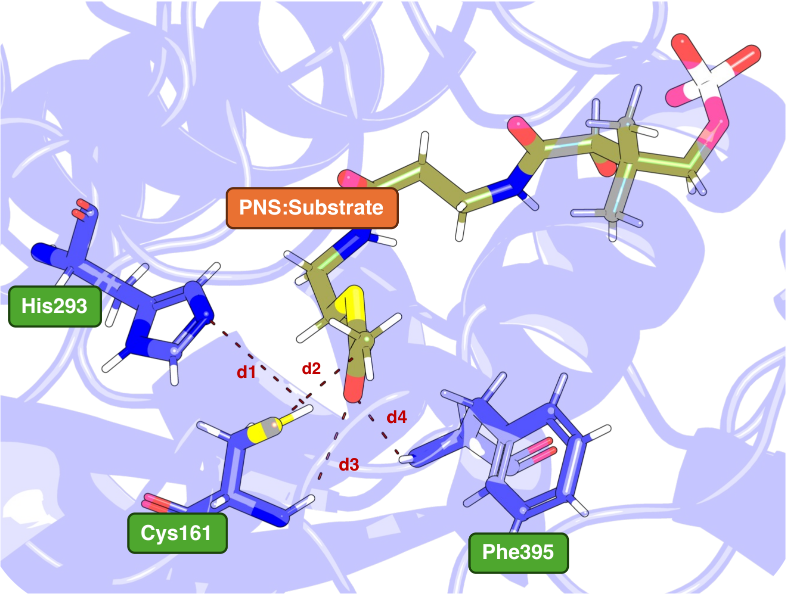


***Figure S17 Visual representation of the four analyzed distances: two catalytic distances and two hydrogen bonds within the oxyanion hole****. The Ppant:Substrate is colored green, and the residues that comprise the reactive site of the KS domain are colored purple. Additionally, each distance is represented by red dashed lines. The figure provides a structural overview highlighting key molecular interactions critical for enzyme function. Distance d1 denotes the proximity between Cys161(SH) and His293(Nε), necessary for thiol deprotonation. Distance d2 illustrates the interaction between Cys161(S) and Ppant(C), indicative of the easiness of the subsequent nucleophilic attack. Distances d3 and d4 correspond to the hydrogen bond lengths of the oxyanion hole, which is crucial for stabilizing the transition state in which the carbonyl oxygen bears a marked negative charge.*

a) Acetyl-CoA

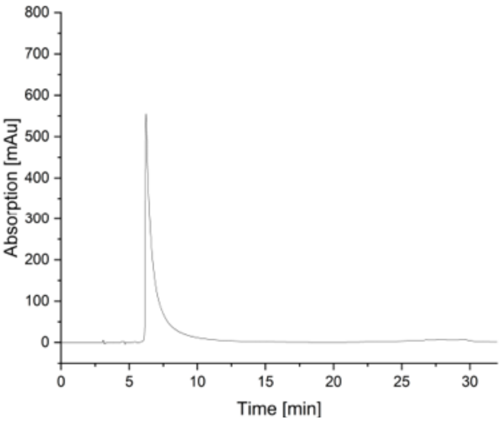


b) hexanoyl-CoA

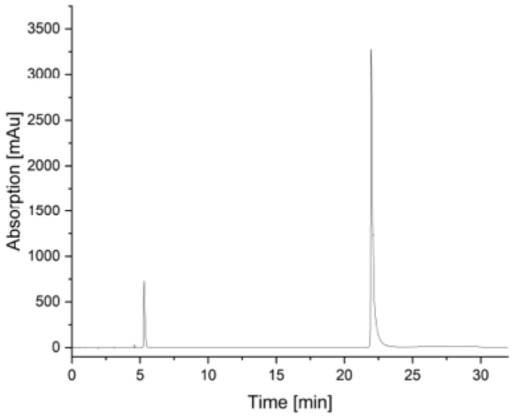


d) octanoyl-CoA

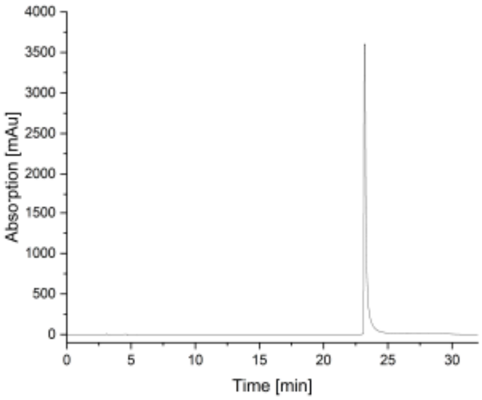


c) decanoyl-CoA

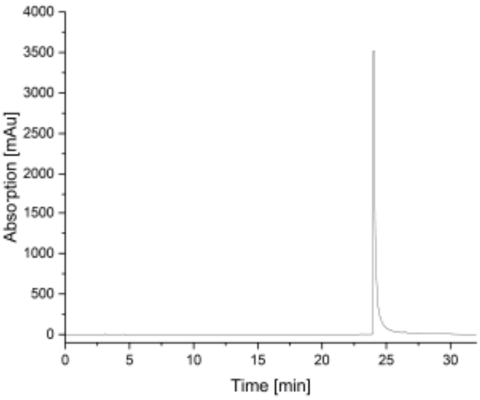

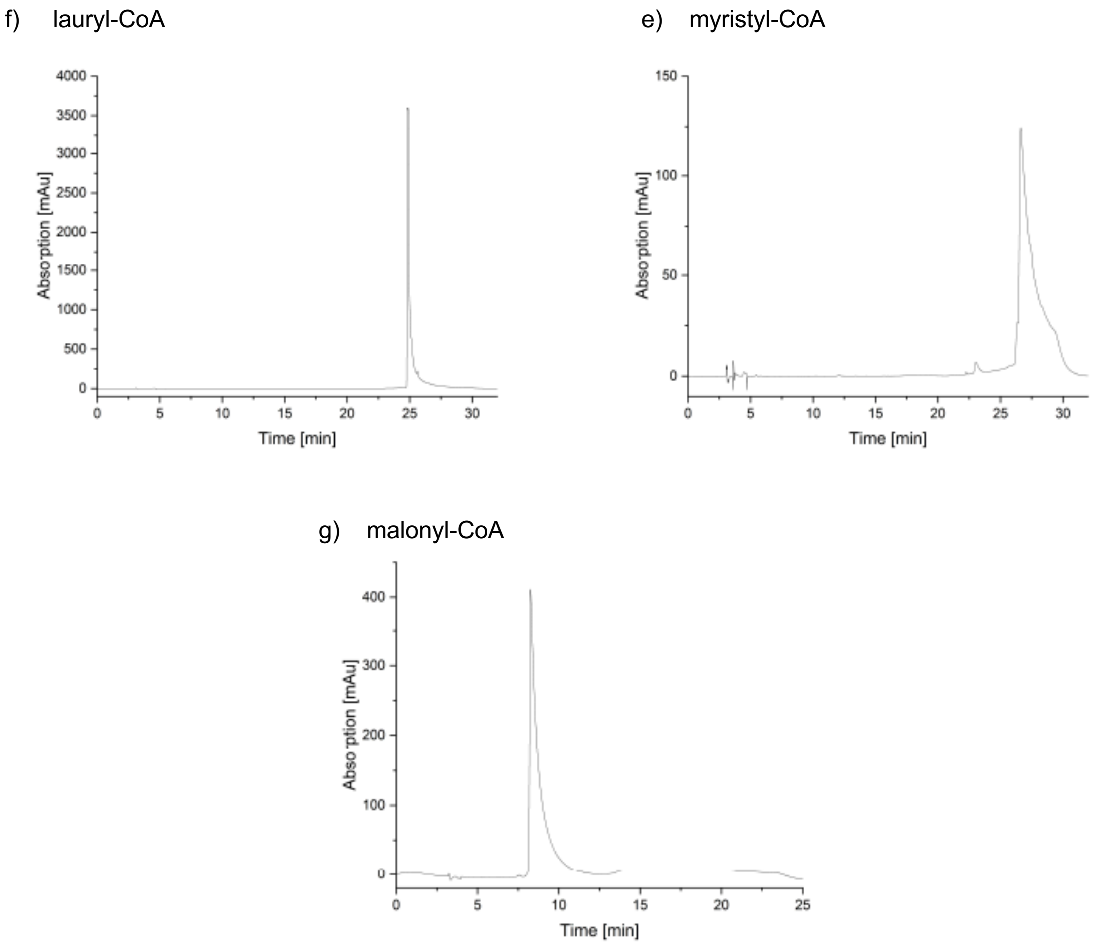


***Figure S18 Representative HPLC chromatograms for acyl-CoA esters.***

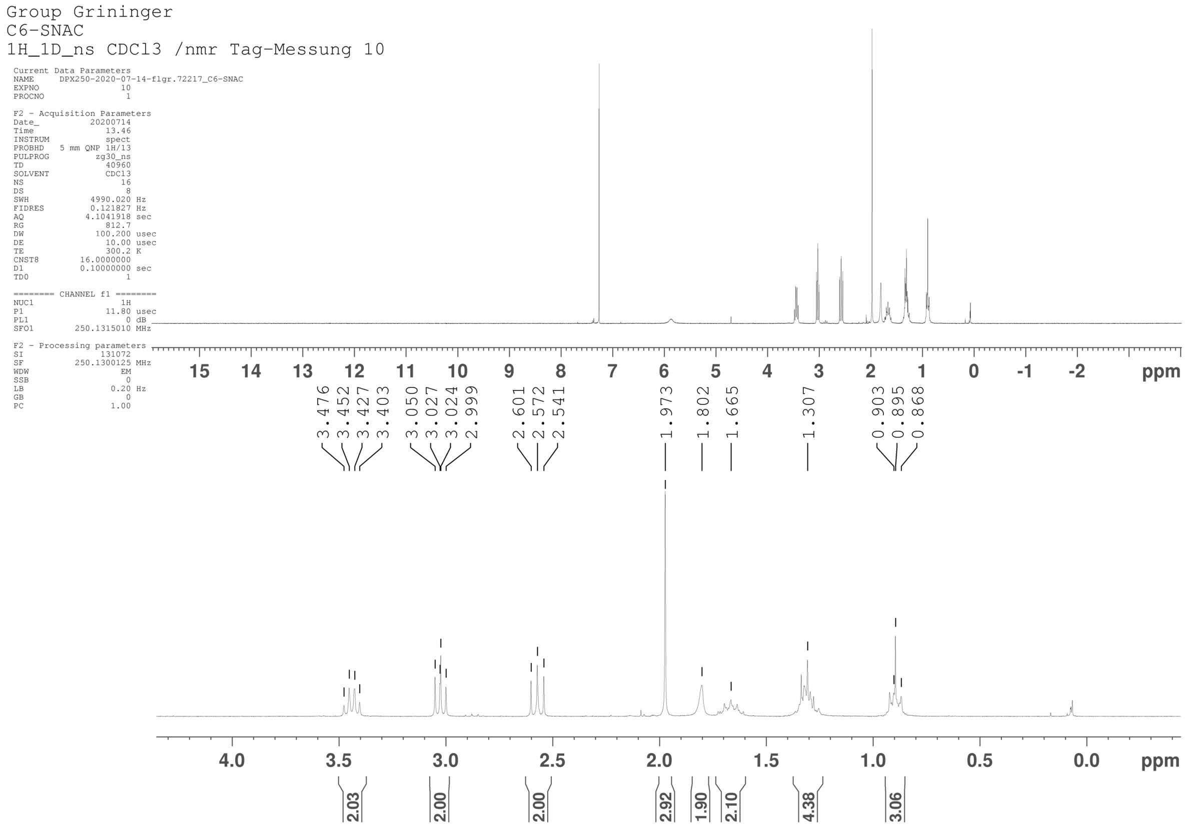


***S19 NMR spectrum of hexanoyl-SNAC**. Top: Spectrum over a wide range. Bottom: Zoom to the range of product signals. (250 MHz, $CDCl_3$)*

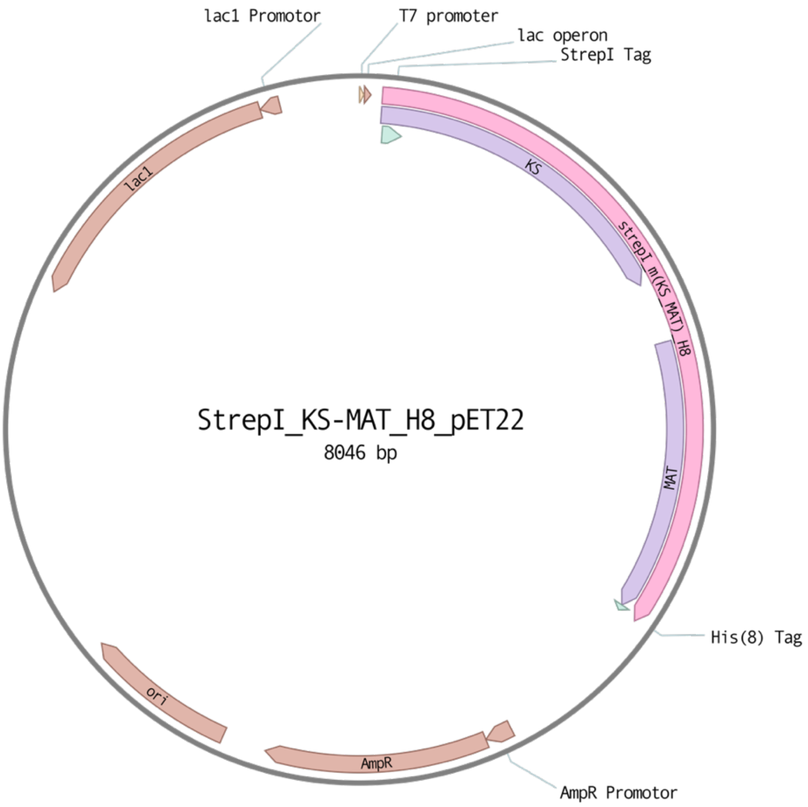


***S20 Plasmid map of KS-MAT**. Vector sites are presented in brown, including lac operon (lac1), origin of replication (ori) and ampicillin resistance (AmpR). The gene encoding for the KS-MAT is shown in pink with the KS and MAT structural part shown in violet respectively. The N-terminal StrepI-Tag and C-terminal 8xHis-Tag are shown in green.*

**List of Sequences:**
**Nucleic acid sequence of template plasmid:**
TAATACGACTCACTATAGGGGAATTGTGAGCGGATAACAATTCCCCTCTAGAAATAATTTTGTTTA
ACTTTAAGAAGGAGATATACATATGAGCGCTTGGAGCCATCCACAATTTGAGAAGGGTGGAGGTT
CTGGCGGTGGATCGGGAGGTTCAGCGTGGAGCCACCCGCAGTTCGAAAAAGGCGCCGGATCCg
aggaggtggtgatagccggtatgtcggggaagttgcccgagtcagagaacctacaggagttctgggccaacctcattggtggtgtggacatgg
tcacagatgatgacaggagatggaaggctgggctctatggattacccaagcggtctggaaagctgaaggatctctccaagttcgacgcctcctt
ttttggggtccaccccaagcaggcacacacaatggacccccagcttcggctgctgttggaagtcagctatgaagcaattgtggatggaggtatc
aacccagcctcactccgaggaacgaacactggcgtctgggtgggtgtgagtggttcagaggcatccgaggcccttagcagagatcccgaga
cgcttctgggctacagcatggtgggctgccagcgtgcaatgatggccaaccggctctctttcttcttcgacttcaaaggaccaagcattgccctgg
acacagcctgctcctccagcttgctggcactacagaatgcctaccaggccatccgtagtggggaatgccccgcggcccttgtgggtgggatca
acctgctcctgaagccgaacacctctgtgcagttcatgaagctgggcatgctcagcccggacggcacctgcagatcctttgatgattcagggag
tggatattgtcgctctgaggctgttgtagcagttctgctgactaagaagtccctggctcggcgggtctatgccacgattctgaatgccggcaccaat
acagatggcagcaaggagcaaggtgtaacattcccctctggagaagtccaagaacaactcatctgctctctgtatcagccagctggtctggcc
ccggagtcgcttgagtatattgaagcccatggcacgggcaccaaggtgggtgacccccaggaactgaatggcattactcggtccctgtgcgcc
ttccgccaggcccctctgttaattggctccaccaaatccaacatgggacaccctgagcctgcctctgggcttgcagccctgaccaaggtgctgtt
atccctggagcatggggtctgggcccctaacctgcacttccacaaccccaaccctgagatcccagcacttcttgatgggcggctgcaggtggtc
gataggcccctgcctgttcgtggtggcaacgtgggcatcaactcatttggcttcggaggctccaatgttcatgtcatcctccagcccaacacacgg
caggcccctgcgcccactgcacacgctgcccttccccatttgctgcacgccagtggacgcaccttagaggcagtgcaggacctgctggaaca
gggccgccagcacagccaggacctggcctttgtgagcatgctcaatgacattgcggcaacccctacagcagccatgcccttcaggggttaca
ctgtgctaggtgttgagggccgtgtccaagaagtgcagcaagtgtccaccaacaagcgcccactctggttcatctgctcagggatgggcacgc
agtggcgcgggatggggctgagcctcatgcgcctggacagcttccgtgagtctatcctgcgctccgatgaggctgtgaagccgttgggagtga
aagtgtcagatctgctgttgagcacagatgagcgcacctttgatgacatcgtgcatgcctttgtgagcctcactgccatccagattgccctcatcga
cctactgacttctgtgggactgaaacctgacggcatcattgggcacGccttgggagaggttgcctgtggctatgcagatggctgtctctcccaga
gagaggctgtgcttgcagcttactggcgaggccagtgcatcaaagatgcccacctcccgcctggatccatggcagctgttggtttgtcctgggag
gaatgtaaacagcgctgccccgctggcgtggtgcctgcctgccacaactctgaggacaccgtgaccatctctggacctcaggctgcagtgaat
gaatttgtggagcagctaaagcaagaaggtgtgtttgccaaggaggtacgaacaggaggcctggctttccactcctacttcatggaaggaattg
cccccacattgctgcaggctctcaagaaggtgatccgggaaccacggccgcgctcggctcgatggctcagcacctctatccctgaggcccagt
ggcagagcagcctggcccgcacatcttctgccgagtacaatgtcaacaacctggtgagccctgtgctcttccaggaagcactgtggcacatcc
ctgagcatgccgtggtgctggagattgcgccccacgcactgttgcaggctgtcctgaagcgaggcgtgaagtccagctgcaccatcattccctt
gatgaagagggatcataaagataacttggagttctttctcaccaaccttggcaaggtgcacctcacaggcatcaatgtcaaccctaacgccttgt
tcccacctgtggagttcccggctccccgagggactcctctcatctcccctcacatcaagtgggaccacagtcagacttgggatgtcccggttgctg
aggacttcccaaacggctcaggttccccctcagccCATCATCACCACCACCACCACCACTGAGATCCGGCTGCTA
ACAAAGCCCGAAAGGAAGCTGAGTTGGCTGCTGCCACCGCTGAGCAATAACTAGCATAACCCCT
TGGGGCCTCTAAACGGGTCTTGAGGGGTTTTTTGCTGAAAGGAGGAACTATATCCGGATTGGCG
AATGGGACGCGCCCTGTAGCGGCGCATTAAGCGCGGCGGGTGTGGTGGTTACGCGCAGCGTGA
CCGCTACACTTGCCAGCGCCCTAGCGCCCGCTCCTTTCGCTTTCTTCCCTTCCTTTCTCGCCACG
TTCGCCGGCTTTCCCCGTCAAGCTCTAAATCGGGGGCTCCCTTTAGGGTTCCGATTTAGTGCTTT
ACGGCACCTCGACCCCAAAAAACTTGATTAGGGTGATGGTTCACGTAGTGGGCCATCGCCCTGA
TAGACGGTTTTTCGCCCTTTGACGTTGGAGTCCACGTTCTTTAATAGTGGACTCTTGTTCCAAACT
GGAACAACACTCAACCCTATCTCGGTCTATTCTTTTGATTTATAAGGGATTTTGCCGATTTCGGCC
TATTGGTTAAAAAATGAGCTGATTTAACAAAAATTTAACGCGAATTTTAACAAAATATTAACGTTTA
CAATTTCAGGTGGCACTTTTCGGGGAAATGTGCGCGGAACCCCTATTTGTTTATTTTTCTAAATAC
ATTCAAATATGTATCCGCTCATGAGACAATAACCCTGATAAATGCTTCAATAATATTGAAAAAGGAA
GAGTATGAGTATTCAACATTTCCGTGTCGCCCTTATTCCCTTTTTTGCGGCATTTTGCCTTCCTGTT
TTTGCTCACCCAGAAACGCTGGTGAAAGTAAAAGATGCTGAAGATCAGTTGGGTGCACGAGTGG
GTTACATCGAACTGGATCTCAACAGCGGTAAGATCCTTGAGAGTTTTCGCCCCGAAGAACGTTTT
CCAATGATGAGCACTTTTAAAGTTCTGCTATGTGGCGCGGTATTATCCCGTATTGACGCCGGGCA
AGAGCAACTCGGTCGCCGCATACACTATTCTCAGAATGACTTGGTTGAGTACTCACCAGTCACAG
AAAAGCATCTTACGGATGGCATGACAGTAAGAGAATTATGCAGTGCTGCCATAACCATGAGTGAT
AACACTGCGGCCAACTTACTTCTGACAACGATCGGAGGACCGAAGGAGCTAACCGCTTTTTTGCA
CAACATGGGGGATCATGTAACTCGCCTTGATCGTTGGGAACCGGAGCTGAATGAAGCCATACCA
AACGACGAGCGTGACACCACGATGCCTGCAGCAATGGCAACAACGTTGCGCAAACTATTAACTG
GCGAACTACTTACTCTAGCTTCCCGGCAACAATTAATAGACTGGATGGAGGCGGATAAAGTTGCA
GGACCACTTCTGCGCTCGGCCCTTCCGGCTGGCTGGTTTATTGCTGATAAATCTGGAGCCGGTG
AGCGTGGGTCTCGCGGTATCATTGCAGCACTGGGGCCAGATGGTAAGCCCTCCCGTATCGTAGT
TATCTACACGACGGGGAGTCAGGCAACTATGGATGAACGAAATAGACAGATCGCTGAGATAGGT
GCCTCACTGATTAAGCATTGGTAACTGTCAGACCAAGTTTACTCATATATACTTTAGATTGATTTAA
AACTTCATTTTTAATTTAAAAGGATCTAGGTGAAGATCCTTTTTGATAATCTCATGACCAAAATCCC
TTAACGTGAGTTTTCGTTCCACTGAGCGTCAGACCCCGTAGAAAAGATCAAAGGATCTTCTTGAG
ATCCTTTTTTTCTGCGCGTAATCTGCTGCTTGCAAACAAAAAAACCACCGCTACCAGCGGTGGTTT

GTTTGCCGGATCAAGAGCTACCAACTCTTTTTCCGAAGGTAACTGGCTTCAGCAGAGCGCAGATA
CCAAATACTGTCCTTCTAGTGTAGCCGTAGTTAGGCCACCACTTCAAGAACTCTGTAGCACCGCC
TACATACCTCGCTCTGCTAATCCTGTTACCAGTGGCTGCTGCCAGTGGCGATAAGTCGTGTCTTA
CCGGGTTGGACTCAAGACGATAGTTACCGGATAAGGCGCAGCGGTCGGGCTGAACGGGGGGTT
CGTGCACACAGCCCAGCTTGGAGCGAACGACCTACACCGAACTGAGATACCTACAGCGTGAGCT
ATGAGAAAGCGCCACGCTTCCCGAAGGGAGAAAGGCGGACAGGTATCCGGTAAGCGGCAGGGT
CGGAACAGGAGAGCGCACGAGGGAGCTTCCAGGGGGAAACGCCTGGTATCTTTATAGTCCTGTC
GGGTTTCGCCACCTCTGACTTGAGCGTCGATTTTTGTGATGCTCGTCAGGGGGGCGGAGCCTAT
GGAAAAACGCCAGCAACGCGGCCTTTTTACGGTTCCTGGCCTTTTGCTGGCCTTTTGCTCACATG
TTCTTTCCTGCGTTATCCCCTGATTCTGTGGATAACCGTATTACCGCCTTTGAGTGAGCTGATACC
GCTCGCCGCAGCCGAACGACCGAGCGCAGCGAGTCAGTGAGCGAGGAAGCGGAAGAGCGCCT
GATGCGGTATTTTCTCCTTACGCATCTGTGCGGTATTTCACACCGCATATATGGTGCACTCTCAGT
ACAATCTGCTCTGATGCCGCATAGTTAAGCCAGTATACACTCCGCTATCGCTACGTGACTGGGTC
ATGGCTGCGCCCCGACACCCGCCAACACCCGCTGACGCGCCCTGACGGGCTTGTCTGCTCCCG
GCATCCGCTTACAGACAAGCTGTGACCGTCTCCGGGAGCTGCATGTGTCAGAGGTTTTCACCGT
CATCACCGAAACGCGCGAGGCAGCTGCGGTAAAGCTCATCAGCGTGGTCGTGAAGCGATTCACA
GATGTCTGCCTGTTCATCCGCGTCCAGCTCGTTGAGTTTCTCCAGAAGCGTTAATGTCTGGCTTC
TGATAAAGCGGGCCATGTTAAGGGCGGTTTTTTCCTGTTTGGTCACTGATGCCTCCGTGTAAGGG
GGATTTCTGTTCATGGGGGTAATGATACCGATGAAACGAGAGAGGATGCTCACGATACGGGTTA
CTGATGATGAACATGCCCGGTTACTGGAACGTTGTGAGGGTAAACAACTGGCGGTATGGATGCG
GCGGGACCAGAGAAAAATCACTCAGGGTCAATGCCAGCGCTTCGTTAATACAGATGTAGGTGTT
CCACAGGGTAGCCAGCAGCATCCTGCGATGCAGATCCGGAACATAATGGTGCAGGGCGCTGAC
TTCCGCGTTTCCAGACTTTACGAAACACGGAAACCGAAGACCATTCATGTTGTTGCTCAGGTCGC
AGACGTTTTGCAGCAGCAGTCGCTTCACGTTCGCTCGCGTATCGGTGATTCATTCTGCTAACCAG
TAAGGCAACCCCGCCAGCCTAGCCGGGTCCTCAACGACAGGAGCACGATCATGCGCACCCGTG
GGGCCGCCATGCCGGCGATAATGGCCTGCTTCTCGCCGAAACGTTTGGTGGCGGGACCAGTGA
CGAAGGCTTGAGCGAGGGCGTGCAAGATTCCGAATACCGCAAGCGACAGGCCGATCATCGTCG
CGCTCCAGCGAAAGCGGTCCTCGCCGAAAATGACCCAGAGCGCTGCCGGCACCTGTCCTACGA
GTTGCATGATAAAGAAGACAGTCATAAGTGCGGCGACGATAGTCATGCCCCGCGCCCACCGGAA
GGAGCTGACTGGGTTGAAGGCTCTCAAGGGCATCGGTCGAGATCCCGGTGCCTAATGAGTGAG
CTAACTTACATTAATTGCGTTGCGCTCACTGCCCGCTTTCCAGTCGGGAAACCTGTCGTGCCAGC
TGCATTAATGAATCGGCCAACGCGCGGGGAGAGGCGGTTTGCGTATTGGGCGCCAGGGTGGTT
TTTCTTTTCACCAGTGAGACGGGCAACAGCTGATTGCCCTTCACCGCCTGGCCCTGAGAGAGTT
GCAGCAAGCGGTCCACGCTGGTTTGCCCCAGCAGGCGAAAATCCTGTTTGATGGTGGTTAACGG
CGGGATATAACATGAGCTGTCTTCGGTATCGTCGTATCCCACTACCGAGATATCCGCACCAACGC
GCAGCCCGGACTCGGTAATGGCGCGCATTGCGCCCAGCGCCATCTGATCGTTGGCAACCAGCA
TCGCAGTGGGAACGATGCCCTCATTCAGCATTTGCATGGTTTGTTGAAAACCGGACATGGCACTC
CAGTCGCCTTCCCGTTCCGCTATCGGCTGAATTTGATTGCGAGTGAGATATTTATGCCAGCCAGC
CAGACGCAGACGCGCCGAGACAGAACTTAATGGGCCCGCTAACAGCGCGATTTGCTGGTGACC
CAATGCGACCAGATGCTCCACGCCCAGTCGCGTACCGTCTTCATGGGAGAAAATAATACTGTTGA
TGGGTGTCTGGTCAGAGACATCAAGAAATAACGCCGGAACATTAGTGCAGGCAGCTTCCACAGC
AATGGCATCCTGGTCATCCAGCGGATAGTTAATGATCAGCCCACTGACGCGTTGCGCGAGAAGA
TTGTGCACCGCCGCTTTACAGGCTTCGACGCCGCTTCGTTCTACCATCGACACCACCACGCTGG
CACCCAGTTGATCGGCGCGAGATTTAATCGCCGCGACAATTTGCGACGGCGCGTGCAGGGCCA
GACTGGAGGTGGCAACGCCAATCAGCAACGACTGTTTGCCCGCCAGTTGTTGTGCCACGCGGTT
GGGAATGTAATTCAGCTCCGCCATCGCCGCTTCCACTTTTTCCCGCGTTTTCGCAGAAACGTGGC
TGGCCTGGTTCACCACGCGGGAAACGGTCTGATAAGAGACACCGGCATACTCTGCGACATCGTA
TAACGTTACTGGTTTCACATTCACCACCCTGAATTGACTCTCTTCCGGGCGCTATCATGCCATACC
GCGAAAGGTTTTGCGCCATTCGATGGTGTCCGGGATCTCGACGCTCTCCCTTATGCGACTCCTG
CATTAGGAAGCAGCCCAGTAGTAGGTTGAGGCCGTTGAGCACCGCCGCCGCAAGGAATGGTGC
ATGCAAGGAGATGGCGCCCAACAGTCCCCCGGCCACGGGGCCTGCCACCATACCCACGCCGAA
ACAAGCGCTCATGAGCCCGAAGTGGCGAGCCCGATCTTCCCCATCGGTGATGTCGGCGATATAG
GCGCCAGCAACCGCACCTGTGGCGCCGGTGATGCCGGCCACGATGCGTCCGGCGTAGAGGAT
CGAGATCTCGATCCCGCGAAAT

**Amino acid sequence of MabA**
MHHHHHHHHTATATEGAKPPFVSRSVLVTGGNRGIGLAIAQRLAADGHKVAVTHRGSGAP
KGLFGVECDVTDSDAVDRAFTAVEEHQGPVEVLVSNAGLSADAFLMRMTEEKFEKVINANL
TGAFRVAQRASRSMQRNKFGRMIFIGSVSGSWGIGNQANYAASKAGVIGMARSIARELSKA
NVTANVVAPGYIDTDMTRALDERIQQGALQFIPAKRVGTPAEVAGVVSFLASEDASYISGAVI
PVDGGMGMGH
The plasmid is based on the pET22 vector and was prepared by Alexander Rittner.

**Amino acid sequence of ACP**
MSAWSHPQFEKGAGDGDTQRDLVKAVAHILGIRDLAGINLDSTLADLGLDSLMGVEVRQILE
REHDLVLPMREVRQLTLRKLQEMSSKTDSATDTTAPLEHHHHHHHH
The plasmid is based on the pET22 vector and was prepared by Alexander Rittner.[42]

**Amino acid sequence of Sfp**
MKIYGIYMDRPLSQEENERFMTFISPEKREKCRRFYHKEDAHRTLLGDVLVRSVISRQYQLD
KSDIRFSTQEYGKPCIPDLPDAHFNISHSGRWVIGAFDSQPIGIDIEKTKPISLEIAKRFFSKTE
YSDLLAKDKDEQTDYFYHLWSMKESFIKQEGKGLSLPLDSFSVRLHQDGQVSIELPDSHSP
CYIKTYEVDPGYKMAVCAAHPDFPEDITMVSYEELLRSHHHHHH
The plasmid is based on the pQE60 vector and was prepared by Peter Tufar.[62]

**Amino acid sequence of KS-MAT$^{S581A}$**
MSAWSHPQFEKGGGSGGGSGGSAWSHPQFEKGAGSEEVVIAGMSGKLPESENLQEFWA
NLIGGVDMVTDDDRRWKAGLYGLPKRSGKLKDLSKFDASFFGVHPKQAHTMDPQLRLLLE
VSYEAIVDGGINPASLRGTNTGVWVGVSGSEASEALSRDPETLLGYSMVGCQRAMMANRL
SFFFDFKGPSIALDTACSSSLLALQNAYQAIRSGECPAALVGGINLLLKPNTSVQFMKLGMLS
PDGTCRSFDDSGSGYCRSEAVVAVLLTKKSLARRVYATILNAGTNTDGSKEQGVTFPSGEV
QEQLICSLYQPAGLAPESLEYIEAHGTGTKVGDPQELNGITRSLCAFRQAPLLIGSTKSNMG
HPEPASGLAALTKVLLSLEHGVWAPNLHFHNPNPEIPALLDGRLQVVDRPLPVRGGNVGINS
FGFGGSNVHVILQPNTRQAPAPTAHAALPHLLHASGRTLEAVQDLLEQGRQHSQDLAFVSM
LNDIAATPTAAMPFRGYTVLGVEGRVQEVQQVSTNKRPLWFICSGMGTQWRGMGLSLMRL
DSFRESILRSDEAVKPLGVKVSDLLLSTDERTFDDIVHAFVSLTAIQIALIDLLTSVGLKPDGIIG
HALGEVACGYADGCLSQREAVLAAYWRGQCIKDAHLPPGSMAAVGLSWEECKQRCPAGV
VPACHNSEDTVTISGPQAAVNEFVEQLKQEGVFAKEVRTGGLAFHSYFMEGIAPTLLQALKK
VIREPRPRSARWLSTSIPEAQWQSSLARTSSAEYNVNNLVSPVLFQEALWHIPEHAVVLEIA
PHALLQAVLKRGVKSSCTIIPLMKRDHKDNLEFFLTNLGKVHLTGINVNPNALFPPVEFPAPR
GTPLISPHIKWDHSQTWDVPVAEDFPNGSGSPSAHHHHHHHH
The plasmid is based on the pET22 vector and was prepared by Alexander Rittner.[24]

**Amino acid sequence of KS$^{C161G}$-MAT$^{S581A}$**
MSAWSHPQFEKGGGSGGGSGGSAWSHPQFEKGAGSEEVVIAGMSGKLPESENLQEFWA
NLIGGVDMVTDDDRRWKAGLYGLPKRSGKLKDLSKFDASFFGVHPKQAHTMDPQLRLLLE
VSYEAIVDGGINPASLRGTNTGVWVGVSGSEASEALSRDPETLLGYSMVGCQRAMMANRL
SFFFDFKGPSIALDTAGSSSLLALQNAYQAIRSGECPAALVGGINLLLKPNTSVQFMKLGMLS
PDGTCRSFDDSGSGYCRSEAVVAVLLTKKSLARRVYATILNAGTNTDGSKEQGVTFPSGEV
QEQLICSLYQPAGLAPESLEYIEAHGTGTKVGDPQELNGITRSLCAFRQAPLLIGSTKSNMG
HPEPASGLAALTKVLLSLEHGVWAPNLHFHNPNPEIPALLDGRLQVVDRPLPVRGGNVGINS
FGFGGSNVHVILQPNTRQAPAPTAHAALPHLLHASGRTLEAVQDLLEQGRQHSQDLAFVSM
LNDIAATPTAAMPFRGYTVLGVEGRVQEVQQVSTNKRPLWFICSGMGTQWRGMGLSLMRL
DSFRESILRSDEAVKPLGVKVSDLLLSTDERTFDDIVHAFVSLTAIQIALIDLLTSVGLKPDGIIG
HALGEVACGYADGCLSQREAVLAAYWRGQCIKDAHLPPGSMAAVGLSWEECKQRCPAGV
VPACHNSEDTVTISGPQAAVNEFVEQLKQEGVFAKEVRTGGLAFHSYFMEGIAPTLLQALKK
VIREPRPRSARWLSTSIPEAQWQSSLARTSSAEYNVNNLVSPVLFQEALWHIPEHAVVLEIA
PHALLQAVLKRGVKSSCTIIPLMKRDHKDNLEFFLTNLGKVHLTGINVNPNALFPPVEFPAPR
GTPLISPHIKWDHSQTWDVPVAEDFPNGSGSPSAHHHHHHHH

**Amino acid sequence of KS$^{D158N}$-MAT$^{S581A}$**

MSAWSHPQFEKGGGSGGGSGGSAWSHPQFEKGAGSEEVVIAGMSGKLPESENLQEFWA
NLIGGVDMVTDDDRRWKAGLYGLPKRSGKLKDLSKFDASFFGVHPKQAHTMDPQLRLLLE
VSYEAIVDGGINPASLRGTNTGVWVGVSGSEASEALSRDPETLLGYSMVGCQRAMMANRL
SFFFDFKGPSIALNTACSSSLLALQNAYQAIRSGECPAALVGGINLLLKPNTSVQFMKLGMLS
PDGTCRSFDDSGSGYCRSEAVVAVLLTKKSLARRVYATILNAGTNTDGSKEQGVTFPSGEV
QEQLICSLYQPAGLAPESLEYIEAHGTGTKVGDPQELNGITRSLCAFRQAPLLIGSTKSNMG
HPEPASGLAALTKVLLSLEHGVWAPNLHFHNPNPEIPALLDGRLQVVDRPLPVRGGNVGINS
FGFGGSNVHVILQPNTRQAPAPTAHAALPHLLHASGRTLEAVQDLLEQGRQHSQDLAFVSM
LNDIAATPTAAMPFRGYTVLGVEGRVQEVQQVSTNKRPLWFICSGMGTQWRGMGLSLMRL
DSFRESILRSDEAVKPLGVKVSDLLLSTDERTFDDIVHAFVSLTAIQIALIDLLTSVGLKPDGIIG
HALGEVACGYADGCLSQREAVLAAYWRGQCIKDAHLPPGSMAAVGLSWEECKQRCPAGV
VPACHNSEDTVTISGPQAAVNEFVEQLKQEGVFAKEVRTGGLAFHSYFMEGIAPTLLQALKK
VIREPRPRSARWLSTSIPEAQWQSSLARTSSAEYNVNNLVSPVLFQEALWHIPEHAVVLEIA
PHALLQAVLKRGVKSSCTIIPLMKRDHKDNLEFFLTNLGKVHLTGINVNPNALFPPVEFPAPR
GTPLISPHIKWDHSQTWDVPVAEDFPNGSGSPSAHHHHHHHH

**Amino acid sequence of KS$^{D158S}$-MAT$^{S581A}$**

MSAWSHPQFEKGGGSGGGSGGSAWSHPQFEKGAGSEEVVIAGMSGKLPESENLQEFWA
NLIGGVDMVTDDDRRWKAGLYGLPKRSGKLKDLSKFDASFFGVHPKQAHTMDPQLRLLLE
VSYEAIVDGGINPASLRGTNTGVWVGVSGSEASEALSRDPETLLGYSMVGCQRAMMANRL
SFFFDFKGPSIALSTACSSSLLALQNAYQAIRSGECPAALVGGINLLLKPNTSVQFMKLGMLS
PDGTCRSFDDSGSGYCRSEAVVAVLLTKKSLARRVYATILNAGTNTDGSKEQGVTFPSGEV
QEQLICSLYQPAGLAPESLEYIEAHGTGTKVGDPQELNGITRSLCAFRQAPLLIGSTKSNMG
HPEPASGLAALTKVLLSLEHGVWAPNLHFHNPNPEIPALLDGRLQVVDRPLPVRGGNVGINS
FGFGGSNVHVILQPNTRQAPAPTAHAALPHLLHASGRTLEAVQDLLEQGRQHSQDLAFVSM
LNDIAATPTAAMPFRGYTVLGVEGRVQEVQQVSTNKRPLWFICSGMGTQWRGMGLSLMRL
DSFRESILRSDEAVKPLGVKVSDLLLSTDERTFDDIVHAFVSLTAIQIALIDLLTSVGLKPDGIIG
HALGEVACGYADGCLSQREAVLAAYWRGQCIKDAHLPPGSMAAVGLSWEECKQRCPAGV
VPACHNSEDTVTISGPQAAVNEFVEQLKQEGVFAKEVRTGGLAFHSYFMEGIAPTLLQALKK
VIREPRPRSARWLSTSIPEAQWQSSLARTSSAEYNVNNLVSPVLFQEALWHIPEHAVVLEIA
PHALLQAVLKRGVKSSCTIIPLMKRDHKDNLEFFLTNLGKVHLTGINVNPNALFPPVEFPAPR
GTPLISPHIKWDHSQTWDVPVAEDFPNGSGSPSAHHHHHHHH

**Amino acid sequence of KS$^{R137K}$-MAT$^{S581A}$**

MSAWSHPQFEKGGGSGGGSGGSAWSHPQFEKGAGSEEVVIAGMSGKLPESENLQEFWA
NLIGGVDMVTDDDRRWKAGLYGLPKRSGKLKDLSKFDASFFGVHPKQAHTMDPQLRLLLE
VSYEAIVDGGINPASLRGTNTGVWVGVSGSEASEALSRDPETLLGYSMVGCQKAMMANRL
SFFFDFKGPSIALDTACSSSLLALQNAYQAIRSGECPAALVGGINLLLKPNTSVQFMKLGMLS
PDGTCRSFDDSGSGYCRSEAVVAVLLTKKSLARRVYATILNAGTNTDGSKEQGVTFPSGEV
QEQLICSLYQPAGLAPESLEYIEAHGTGTKVGDPQELNGITRSLCAFRQAPLLIGSTKSNMG
HPEPASGLAALTKVLLSLEHGVWAPNLHFHNPNPEIPALLDGRLQVVDRPLPVRGGNVGINS
FGFGGSNVHVILQPNTRQAPAPTAHAALPHLLHASGRTLEAVQDLLEQGRQHSQDLAFVSM
LNDIAATPTAAMPFRGYTVLGVEGRVQEVQQVSTNKRPLWFICSGMGTQWRGMGLSLMRL
DSFRESILRSDEAVKPLGVKVSDLLLSTDERTFDDIVHAFVSLTAIQIALIDLLTSVGLKPDGIIG
HALGEVACGYADGCLSQREAVLAAYWRGQCIKDAHLPPGSMAAVGLSWEECKQRCPAGV
VPACHNSEDTVTISGPQAAVNEFVEQLKQEGVFAKEVRTGGLAFHSYFMEGIAPTLLQALKK
VIREPRPRSARWLSTSIPEAQWQSSLARTSSAEYNVNNLVSPVLFQEALWHIPEHAVVLEIA
PHALLQAVLKRGVKSSCTIIPLMKRDHKDNLEFFLTNLGKVHLTGINVNPNALFPPVEFPAPR
GTPLISPHIKWDHSQTWDVPVAEDFPNGSGSPSAHHHHHHHH

**Amino acid sequence of KS$^{R137A}$-MAT$^{S581A}$**

MSAWSHPQFEKGGGSGGGSGGSAWSHPQFEKGAGSEEVVIAGMSGKLPESENLQEFWA
NLIGGVDMVTDDDRRWKAGLYGLPKRSGKLKDLSKFDASFFGVHPKQAHTMDPQLRLLLE
VSYEAIVDGGINPASLRGTNTGVWVGVSGSEASEALSRDPETLLGYSMVGCQAAMMANRL
SFFFDFKGPSIALDTACSSSLLALQNAYQAIRSGECPAALVGGINLLLKPNTSVQFMKLGMLS
PDGTCRSFDDSGSGYCRSEAVVAVLLTKKSLARRVYATILNAGTNTDGSKEQGVTFPSGEV
QEQLICSLYQPAGLAPESLEYIEAHGTGTKVGDPQELNGITRSLCAFRQAPLLIGSTKSNMG
HPEPASGLAALTKVLLSLEHGVWAPNLHFHNPNPEIPALLDGRLQVVDRPLPVRGGNVGINS
FGFGGSNVHVILQPNTRQAPAPTAHAALPHLLHASGRTLEAVQDLLEQGRQHSQDLAFVSM
LNDIAATPTAAMPFRGYTVLGVEGRVQEVQQVSTNKRPLWFICSGMGTQWRGMGLSLMRL
DSFRESILRSDEAVKPLGVKVSDLLLSTDERTFDDIVHAFVSLTAIQIALIDLLTSVGLKPDGIIG
HALGEVACGYADGCLSQREAVLAAYWRGQCIKDAHLPPGSMAAVGLSWEECKQRCPAGV
VPACHNSEDTVTISGPQAAVNEFVEQLKQEGVFAKEVRTGGLAFHSYFMEGIAPTLLQALKK
VIREPRPRSARWLSTSIPEAQWQSSLARTSSAEYNVNNLVSPVLFQEALWHIPEHAVVLEIA
PHALLQAVLKRGVKSSCTIIPLMKRDHKDNLEFFLTNLGKVHLTGINVNPNALFPPVEFPAPR
GTPLISPHIKWDHSQTWDVPVAEDFPNGSGSPSAHHHHHHHH

**Amino acid sequence of KS$^{A160G}$-MAT$^{S581A}$**

MSAWSHPQFEKGGGSGGGSGGSAWSHPQFEKGAGSEEVVIAGMSGKLPESENLQEFWA
NLIGGVDMVTDDDRRWKAGLYGLPKRSGKLKDLSKFDASFFGVHPKQAHTMDPQLRLLLE
VSYEAIVDGGINPASLRGTNTGVWVGVSGSEASEALSRDPETLLGYSMVGCQRAMMANRL
SFFFDFKGPSIALDTGCSSSLLALQNAYQAIRSGECPAALVGGINLLLKPNTSVQFMKLGMLS
PDGTCRSFDDSGSGYCRSEAVVAVLLTKKSLARRVYATILNAGTNTDGSKEQGVTFPSGEV
QEQLICSLYQPAGLAPESLEYIEAHGTGTKVGDPQELNGITRSLCAFRQAPLLIGSTKSNMG
HPEPASGLAALTKVLLSLEHGVWAPNLHFHNPNPEIPALLDGRLQVVDRPLPVRGGNVGINS
FGFGGSNVHVILQPNTRQAPAPTAHAALPHLLHASGRTLEAVQDLLEQGRQHSQDLAFVSM
LNDIAATPTAAMPFRGYTVLGVEGRVQEVQQVSTNKRPLWFICSGMGTQWRGMGLSLMRL
DSFRESILRSDEAVKPLGVKVSDLLLSTDERTFDDIVHAFVSLTAIQIALIDLLTSVGLKPDGIIG
HALGEVACGYADGCLSQREAVLAAYWRGQCIKDAHLPPGSMAAVGLSWEECKQRCPAGV
VPACHNSEDTVTISGPQAAVNEFVEQLKQEGVFAKEVRTGGLAFHSYFMEGIAPTLLQALKK
VIREPRPRSARWLSTSIPEAQWQSSLARTSSAEYNVNNLVSPVLFQEALWHIPEHAVVLEIA
PHALLQAVLKRGVKSSCTIIPLMKRDHKDNLEFFLTNLGKVHLTGINVNPNALFPPVEFPAPR
GTPLISPHIKWDHSQTWDVPVAEDFPNGSGSPSAHHHHHHHH

**Amino acid sequence of KS$^{A160V}$-MAT$^{S581A}$**

MSAWSHPQFEKGGGSGGGSGGSAWSHPQFEKGAGSEEVVIAGMSGKLPESENLQEFWA
NLIGGVDMVTDDDRRWKAGLYGLPKRSGKLKDLSKFDASFFGVHPKQAHTMDPQLRLLLE
VSYEAIVDGGINPASLRGTNTGVWVGVSGSEASEALSRDPETLLGYSMVGCQRAMMANRL
SFFFDFKGPSIALDTVCSSSLLALQNAYQAIRSGECPAALVGGINLLLKPNTSVQFMKLGMLS
PDGTCRSFDDSGSGYCRSEAVVAVLLTKKSLARRVYATILNAGTNTDGSKEQGVTFPSGEV
QEQLICSLYQPAGLAPESLEYIEAHGTGTKVGDPQELNGITRSLCAFRQAPLLIGSTKSNMG
HPEPASGLAALTKVLLSLEHGVWAPNLHFHNPNPEIPALLDGRLQVVDRPLPVRGGNVGINS
FGFGGSNVHVILQPNTRQAPAPTAHAALPHLLHASGRTLEAVQDLLEQGRQHSQDLAFVSM
LNDIAATPTAAMPFRGYTVLGVEGRVQEVQQVSTNKRPLWFICSGMGTQWRGMGLSLMRL
DSFRESILRSDEAVKPLGVKVSDLLLSTDERTFDDIVHAFVSLTAIQIALIDLLTSVGLKPDGIIG
HALGEVACGYADGCLSQREAVLAAYWRGQCIKDAHLPPGSMAAVGLSWEECKQRCPAGV
VPACHNSEDTVTISGPQAAVNEFVEQLKQEGVFAKEVRTGGLAFHSYFMEGIAPTLLQALKK
VIREPRPRSARWLSTSIPEAQWQSSLARTSSAEYNVNNLVSPVLFQEALWHIPEHAVVLEIA
PHALLQAVLKRGVKSSCTIIPLMKRDHKDNLEFFLTNLGKVHLTGINVNPNALFPPVEFPAPR
GTPLISPHIKWDHSQTWDVPVAEDFPNGSGSPSAHHHHHHHH

##### List of primers

| | |
|---|---|
| prCG048_KS_D158N_f | ccaagcattgccctgaacacagcctgctcctccag |
| prCG050_KS_D158S_f | ccaagcattgccctgtccacagcctgctcctccag |
| prCG051_KS_R137K_f | atggtgggctgccagaaagcaatgatggccaaccggc |
| prCG053_KS_R137A_f | atggtgggctgccaggcggcaatgatggccaaccggc |
| prCG054_KS_A160G_f | cattgccctggacacaggctgctcctccagcttgctgg |
| prCG056_KS_A160V_f | cattgccctggacacagtctgctcctccagcttgctgg |
| prCG057_KS_D158S_r | ctggaggagcaggctgtGGAcagggcaatgcttgg |
| prCG058_KS_D158N_r | ctggaggagcaggctgtgtTcagggcaatgcttgg |
| prCG059_KS_R137K_r | gccggttggccatcattgcTTTctggcagcccaccat |
| prCG060_KS_R137A_r | gccggttggccatcattgcCGCctggcagcccaccat |
| prCG061_KS_A160G_r | ccagcaagctggaggagcaGCCtgtgtccagggcaatg |
| prCG062_KS_A160V_r | ccagcaagctggaggagcaGACtgtgtccagggcaatg |
